# Autophagy inhibition induces AML cell death and enhances the efficacy of chemotherapy under hypoxia

**DOI:** 10.1101/2024.11.24.625107

**Authors:** Hannah R.S. Fay, Kaitlyn M. Dykstra, Michael N. Phelps, Matthew Johnson, Annemarie Harrigan, Marianna C. Giorgi, Eunice S. Wang

## Abstract

Outcomes of relapsed/refractory acute myeloid leukemia (AML) are poor, and strategies to improve outcomes are urgently needed. One important factor promoting relapse and chemoresistance is the ability of AML cells to thrive *in vivo* within an intrinsically hypoxic bone marrow microenvironment. Here we show that human AML cells exhibit enhanced autophagy, specifically mitophagy (i.e., increased accumulation of mitochondria and decreased mitochondrial membrane potential) under hypoxia. To target this pathway, we investigated the activity of the potent chloroquine-derived autophagy inhibitor, Lys05, on human AML cells, patient samples, and patient derived xenograft models. Inhibition of autophagy by Lys05 in AML cells prevented removal of damaged mitochondria and preferentially enhanced cell death under hypoxia mirroring the marrow microenvironment. Lys05 eradicated human AML cells of all genotypes including p53 mutant cells. Lys05 treatment in primary AML xenografted mice decreased CD34+CD38- human cells and prolonged overall survival. Moreover, Lys05 overcame hypoxia-induced chemoresistance and improved the efficacy of cytarabine, venetoclax, and azacytidine *in vitro* and *in vivo* in AML models. Our results demonstrate the importance of autophagy, specifically mitophagy, as a critical survival and chemoresistance mechanism of AML cells under hypoxic marrow conditions. Therapeutic targeting of this pathway in future clinical studies for AML is warranted.

## INTRODUCTION

Acute myeloid leukemia (AML) is an aggressive hematological malignancy arising from hematopoietic stem and progenitor cells in the bone marrow (BM).^1^ AML has a dismal five-year overall survival rate of 30-35%, largely due to high rates of relapse.^1,2^ While several targeted therapies have been approved for AML, these agents are active only in specific molecular subtypes constituting < 50% of newly diagnosed patients.^2-4^ Moreover, relapse following upfront therapy remains the major clinical challenge limiting long-term survival.^4-6^ Identification of the underlying mechanisms promoting non-genetic chemoresistance in AML cells and development of novel therapeutic strategies to overcome these are urgently needed to improve outcomes for the majority of patients.^7^

The hypoxic bone marrow microenvironment (BMME) is known to play a critical role in mediating therapy resistance in acute leukemia cells.^6,8-11^ AML cells originate and primarily reside in the bone marrow microenvironment, predominantly within specialized marrow niches characterized by intrinsically low oxygenation levels ranging from 0.6% – 2.8%. Further reductions in oxygen levels may occur in the context of rapid disease expansion within the BMME, suggesting that AML cells either specifically adapt to or are preferentially selected to survive under acute and chronic hypoxia.^11-13^ Of note, many conventional cytotoxic chemotherapy agents which target rapidly proliferating cells are markedly less efficacious against AML cells under hypoxic conditions where cancer cells are less metabolically active or even quiescent.^9,14^ In addition, experimental data has demonstrated that the culturing of AML cells under chronic hypoxic conditions mimicking the hypoxic BMME can also induce key survival pathways that in turn confer additional resistance to chemotherapy.^9,11,14^ Despite its importance, to date few studies have investigated the impact of the hypoxic microenvironment on AML chemoresistance and/or explored therapeutic approaches to overcome this scenario in clinically relevant models.

Autophagy is a complex pro-survival recycling process which reallocates nutrients and removes damaged organelles in response to a variety of stressors including starvation and human disease.^15^ Autophagy is known to play an important role in AML development, progression, and resistance.^16^ Prior studies have demonstrated that autophagy promotes leukemic transformation of normal hematopoietic stem cells and supports leukemia stem cell (LSC) functions as well as serving as an induced resistance mechanism.^17-20^ Numerous studies have demonstrated that interrogation of autophagy via genetic knockdown models of autophagy genes or pharmacologic inhibitors acting at different stages of the autophagic pathway can limit AML progression.^21-28^ In our prior work, we demonstrated that human AML cells markedly upregulate autophagy under low oxygen conditions mirroring those of the BMME. Moreover, we showed that autophagy inhibition via siRNA or the pharmacologic agent, chloroquine (CQ) in AML cells under these conditions results in cell death.

Here we identify mitophagy as a key survival and resistance property of human AML cells under chronic hypoxia mimicking the BMME. We demonstrate that mitophagy can be effectively and specifically targeted by the CQ derivative, Lys05, to induce AML cell death, preferentially under hypoxic conditions mimicking the marrow microenvironment. We further demonstrate the broad anti-leukemic activity of Lys05 and the ability of Lys05 to overcome hypoxia-induced chemoresistance to multiple agents *in vitro* and *in vivo* in human AML xenograft and serially transplanted patient-derived xenograft models. These studies provide the rationale for the future clinical targeting of mitophagy for AML therapy, particularly in combinatorial regimens.

## METHODS

### Cells and Patient Samples

Human AML cell lines were obtained from ATCC and DSMZ. Cells were cultured in supplemented RPMI-1640 or IMDM medium. HEL cells were stably transfected as described previously^29^ to generate HEL-luc cell line. Stably transduced MOLM13-GFP-Luc cells (MOLM13-BLIV^30^) were provided by Monica Guzman (Weill Cornell Medicine, NY). Cells were cultured under 21% O_2_ for normoxia or 1% O_2_ for hypoxia using a ProOx-C21 Hypoxia Subchamber (BioSpherix). Cryopreserved ficolled mononuclear cells from de-identified patients with confirmed AML diagnoses were obtained from institute biorepositories (RPCCC, University of Rochester). All individuals provided informed consent under Institutional Review Board approved protocols. Short-term cultures of patient BM cells were performed as previously described.^31^ Characteristics of samples utilized are found in Supplemental Data Table.

### In Vitro Assays

Cells were treated with Lys05 (Selleck Chemicals), chloroquine (Sigma-Aldrich), cytarabine (institutional clinical supply), azacytidine (Selleck), doxorubicin (Selleck), and venetoclax (MedChemExpress). Cell proliferation was measured by WST-8 assay (Cell Counting Kit-8, Dojindo). Synergy analysis was conducted using CompuSyn software (ComboSyn Inc.) and combination index values were calculated by the Chou-Talalay method.^32^ Flow cytometric analysis was performed on a LSRII Flow Cytometer (BD Biosciences) following cell staining with Annexin V-FITC/Propidium Iodide (Apoptosis Detection Kit, BD Pharmingen) or CytoID (CYTO-ID Autophagy Detection Kit, Enzo). Colony formation assays were established using primary AML BM cells cultured at 1×10^5^ cells/ml in MethoCult-H4434 (Stem Cell Technologies) in the presence of vehicle or drug. Colonies were counted on day 13 using light microscopy.

### Immunoblotting

Western blot analysis utilized primary antibodies [anti-LC3B, anti-p62/SQSTM1, anti-PINK1, anti-BNIP3 antibodies (Cell Signaling Technology), anti-Actin (Santa Cruz Biotechnology)]. Detection utilized HRP-linked secondary antibodies (anti-mouse/anti-rabbit (Table 7.5) and chemiluminescence (ECL plus, GE Healthcare, Pittsburgh, PA). Results were acquired using a LI-COR Imager and analyzed with Image Studio Lite (LI-COR).

### Confocal Microscopy

Cells were spun onto poly-L-lysine (Sigma Aldrich)–coated coverslips, washed with PBS, and incubated with Cyto-ID (1:100; Enzo Life Sciences) and MitoTracker Deep Red (ThermoFisher) in PBS with 15% fetal bovine serum at 37°C for 30 minutes in the dark. Coverslips were washed, fixed with 4% paraformaldehyde and mounted with VectaShield with DAPI (49,6-diamidino-2-phenylindole, Vector Laboratories). Microscopy was performed with a Leica TCS SP2 spectral confocal microscope with a 633 objective at the Flow and Image Cytometry Core Facility at Roswell Park. Images were processed by ImageJ software (NIH).

### Murine xenograft models

Systemic bioluminescent human AML xenografts were established using MOLM13-BLIV cells (2.5×10^5^ cells/mouse) inoculated via tail vein injection into NSG mice (Jackson Laboratories or RPCCC). Animals were monitored weekly for leukemic burden by serial bioluminescent imaging (BLI) using Xenogen IVIS50 System (Caliper Life Science) following intraperitoneal (i.p.) D-luciferin injection as previously described.^13^ Animals were treated with vehicle, Lys05 (40mg/kg i.p. for 18 days), CQ (40mg/kg i.p. for 18 days), cytarabine (40mg/kg i.p. for 5 days), venetoclax (100mg/kg p.o. for 5 days/week for 3 weeks), azacitidine (2.5mg/kg i.p. for 5 days) or in combination at the same doses and schedule. PDX models were established following tail vein injection of AML patient BM cells (5-10×10^6^ cells/mouse) into NSG mice. Engraftment (defined as ≥0.1% huCD45^+^muCD45^-^cells in the BM ^33^) was monitored by femoral BM aspirates followed by flow cytometric analysis (BD LSRII Flow Cytometer or Cytek Aurora Spectral Flow Cytometer; FCS Express software) using anti-mouse CD45-BV605, anti-human CD45-PE-Cy5, CD34-PE, and CD38-APC antibodies (BioLegend). Treatment was vehicle or Lys05 (40mg/kg i.p. for 18 days). BM was harvested on day 19 for flow cytometry.

### Statistics

Statistical evaluation of differences in mean values was conducted using two-tailed unpaired student’s t-tests with unequal variance. Overall survival analysis was conducted using log-rank (Mantel-Cox) tests. Error bars represent ± standard deviation. Analysis conducted with GraphPad Prism Software version 8, significance level of p<0.05. In figures, *= p≤0.05, **= p≤0.01, ***= p≤0.001, ****= p≤0.0001, ns= no significance.

## RESULTS

### Lys05 exerts more potent autophagy inhibition than chloroquine in AML

We first compared the potency of autophagy inhibition and anti-leukemic activity mediated by Lys05 to its parent compound, CQ, in three human AML cell lines with diverse molecular profiles: MOLM13 (*KMT2A*-rearranged, *FLT3-ITD* mutant), HL60 (*NRAS* mutation), HEL (*p53* and *JAK2-V617F* mutant). Lys05 acts by preventing autophagosome-lysosome fusion and thus degradation,^34^ therefore treatment should result in increased LC3-II levels. Lys05 treatment resulted in an 8- to 12-fold higher LC3-II levels than control and >7-fold higher levels than CQ-treated cells (**Figure 1A; Supplemental Figure 1A-B**). As LC3-II can be induced by either an increase or block in autophagic flux, we also measured p62 which accumulates with reduced flux and discriminates between the two possible sources of increased LC3-II. Consistent with inhibition of autophagic flux, p62 levels increased in AML cells after Lys05 treatment. We confirmed the enhanced potency of Lys05 over CQ using CytoID, a dye that labels autophagic vesicles and allows for quantification of autophagy via flow cytometry (**Figure 1B**). Lys05 significantly reduced AML cell viability (**Figure 1C; Supplemental Figure 1C-D, G**) and induced higher levels of apoptosis than CQ (74% vs. <5%, p=0.0034) in all three cell lines (**Figure 1D; Supplemental Figure 1E-F**). We further compared the anti-leukemic efficacy of Lys05 and CQ treatment in NSG mice systemically engrafted with a luciferase-expressing human AML cell line (MOLM13-BLIV). After 15 days, Lys05 treatment had significantly reduced leukemia burden compared to vehicle-treated mice (p=0.0023) while there was no difference in leukemia burden between CQ- and vehicle-treated mice (**Figure 1E-F**). Notably, Lys05 significantly prolonged survival over vehicle (p=0.0123) and CQ (p=0.0031; **Figure 1G**).

**Figure 1.**
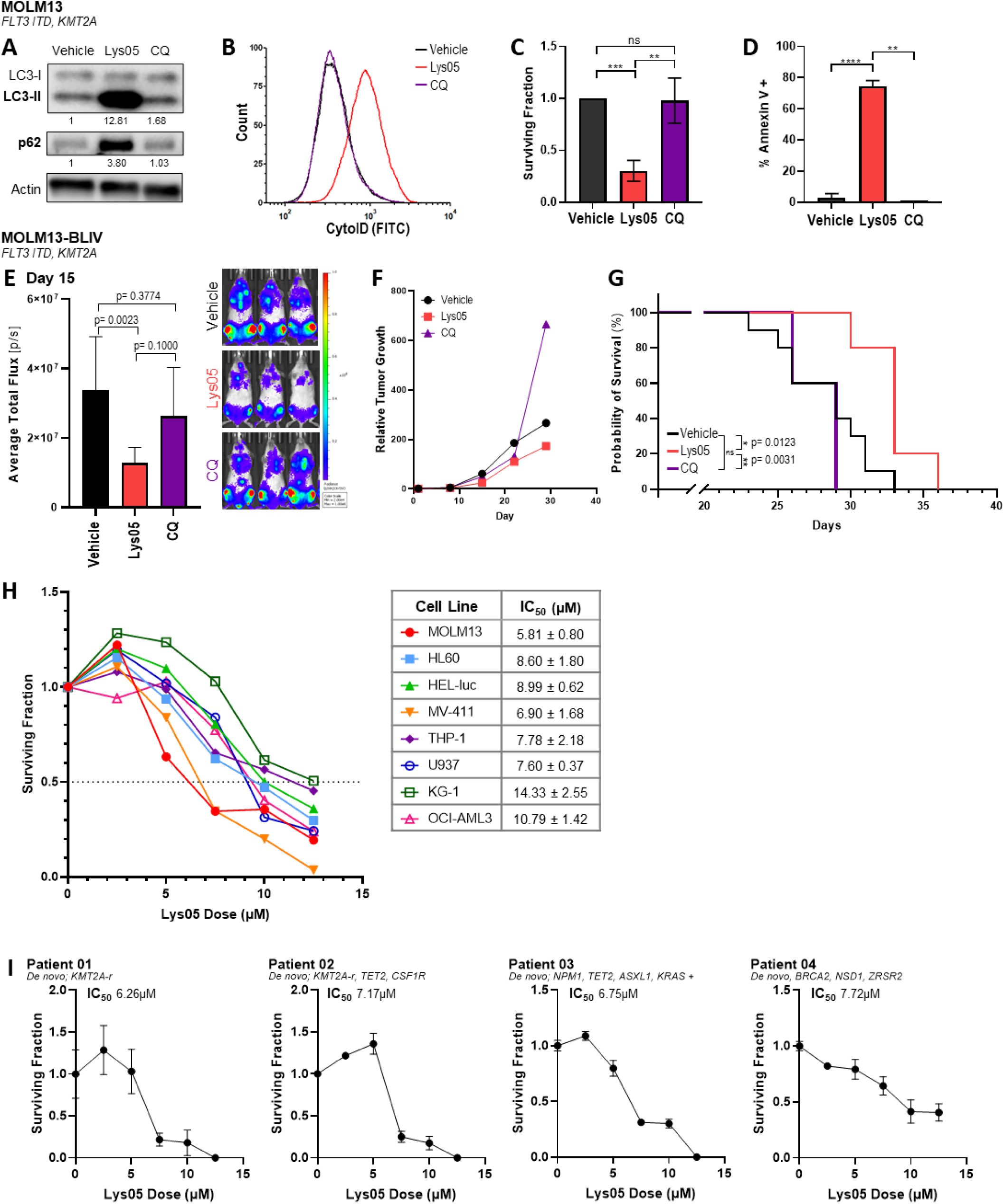
Lys05 exerts greater anti-leukemic activity than chloroquine and decreases survival across multiple AML mutational profiles. (A-D) Human AML cell line MOLM13 was treated with vehicle (PBS), 10µM Lys05, or 10µM CQ for 48 hours, unless otherwise noted. (A) Immunoblot blot analysis of autophagy markers LC3-II and p62 shown along with actin loading control. Quantification shown below blot – each marker was normalized to actin then normalized to vehicle. (B) CytoID flow cytometry for autophagy levels following 72-hour treatment. (C) WST-8 assay for viability. (D) Annexin V-FITC/PI assay for apoptosis. (E-G) NSG mice were systemically engrafted with luciferase-expressing MOLM13-BLIV human AML cells and treated i.p. with vehicle (PBS), Lys05 (40mg/kg), or CQ (40mg/kg) for 18 days. n=5 mice per group. Tumor burden was assessed by BLI. (E) AML tumor burden on day 15 of treatment. Quantification of bioluminescence (left) with representative BLI images shown (right). (F) Relative leukemia burden over time (each mouse was normalized to its Day 1 pre-treatment BLI value). (E) Overall survival. (H) Human AML cell lines were treated with Lys05 for 48 hours followed by WST-8 assay for viability. Left: representative data for each cell line; Right: average IC_50_ values (mean ± standard deviation). Individual curves for each cell line are found in Supplemental Figure 3. (H) Primary AML patient bone marrow samples were treated with Lys05 for 48 hours followed by WST-8 assay for viability. Mutational profile of each cell line or patient sample shown in italics. (C,D) Average of 3 independent experiments shown. (A,B,H) Representative data of 3-4 independent experiments shown. (I) Data from single experiment shown due to limited availability of primary samples. BLI= bioluminescent imaging; CQ=chloroquine.

### Lys05 induces cell death in AML cells with diverse mutational profiles

We evaluated the anti-leukemic activity of Lys05 in eight human AML cell lines reflecting the spectrum of biologically distinct AML subtypes (**Table 1**). While there was some variation, human AML cell lines displayed universal sensitivity to Lys05 treatment, demonstrated by dose-dependent decreases in cell viability and 50% inhibitory concentrations (IC_50_) ranging from 5.8–14.3µM, averaging 8.85µM (**Figure 1H**). Next, we evaluated the ability of Lys05 to inhibit growth of four primary AML patient BM samples with diverse cytogenetic and mutational profiles (**Supplemental Table 1**). Lys05 treatment of primary AML cells cultured *in vitro* short-term consistently resulted in dose-dependent growth inhibition with IC_50_ ranging from 6.3-7.2µM, corroborating our cell line data (**Figure 1I**).

**Table 1.** Comparison of Lys05 IC_50_ values in human AML cell lines under normoxic and hypoxic conditions. Average Lys05 IC_50_ values (mean ± standard deviation, n=3-4 per cell line) for AML cell lines under normoxic (21% O_2_) or hypoxic (1% O_2_) conditions along with mutational status of each cell line. P values represent unpaired t-test between normoxia and hypoxia average IC_50_ values for each cell line, n=3-4 per cell line. Sig= significance.

| Cell Line | Lys05 IC<sub>50</sub> (μM) |  |  |  | Mutational Status |  |  |  |  |  |  |
| --- | --- | --- | --- | --- | --- | --- | --- | --- | --- | --- | --- |
|  | Normoxia<br>(21% O<sub>2</sub>) | Hypoxia<br>(1% O<sub>2</sub>) | P value | Sig | FLT3<br>ITD | TP53 | KMT2A | NRAS | PTEN | DNMT<br>3A | NPM1 |
| **MOLM13** | 5.81 ± 0.80 | 4.13 ± 0.41 | 0.0316 | * | X |  | X |  |  |  |  |
| **HL60** | 8.60 ± 1.80 | 4.62 ± 1.28 | 0.0051 | ** |  | Null |  | X |  |  |  |
| **HEL-luc** | 8.99 ± 0.62 | 5.99 ± 1.27 | 0.0055 | ** |  | X |  |  |  |  |  |
| **MV411** | 6.90 ± 1.68 | 4.08 ± 1.00 | 0.0663 | ns | X |  | X |  |  |  |  |
| **THP-1** | 7.78 ± 2.18 | 4.31 ± 0.93 | 0.0648 | ns |  | X |  | X |  |  |  |
| **U937** | 7.60 ± 0.37 | 4.53 ± 0.18 | 0.0000058 | **** |  | X |  |  | X |  |  |
| **KG-1** | 14.33 ± 2.55 | 7.61 ± 1.33 | 0.0154 | * |  | X |  | X |  |  |  |
| **OCI-AML3** | 10.79 ± 1.42 | 5.98 ± 1.63 | 0.0043 | ** |  |  |  | X |  | X | X |

### Lys05 exerts enhanced anti-leukemic effects under hypoxia

Our prior work demonstrated that AML cells exhibit increased dependence on autophagy under hypoxia and are subsequently more sensitive to autophagy inhibition under low oxygen conditions mirroring the BMME.^35^ Therefore, we compared the ability of Lys05 to inhibit autophagy and exert anti-proliferative effects in human AML cells cultured under hypoxic (1% O_2_) vs. normoxic (21% O_2_) conditions. The basal LC3-II level was higher under hypoxia and Lys05 induced a further increase in LC3-II levels under hypoxia compared to normoxia in all three cell lines (**Figure 2A, Supplemental Figure 2A-B**). The basal level of p62 was decreased under hypoxia, indicating increased autophagic flux under hypoxic conditions, however Lys05 treatment caused p62 levels to increase, indicative of the block in autophagic flux. The enhanced activity of Lys05 under hypoxia was confirmed by CytoID flow cytometry (**Figure 2B, Supplemental Figure 2C-D**). Lys05 caused a dose-dependent increase in CytoID in all three cell lines, with further enhancement under hypoxia (**Figure 2B, Supplemental Figure 2C-D**), suggesting that the enhanced activity of Lys05 under hypoxia is due to increased levels of autophagy occurring in AML cells.

**Figure 2.**
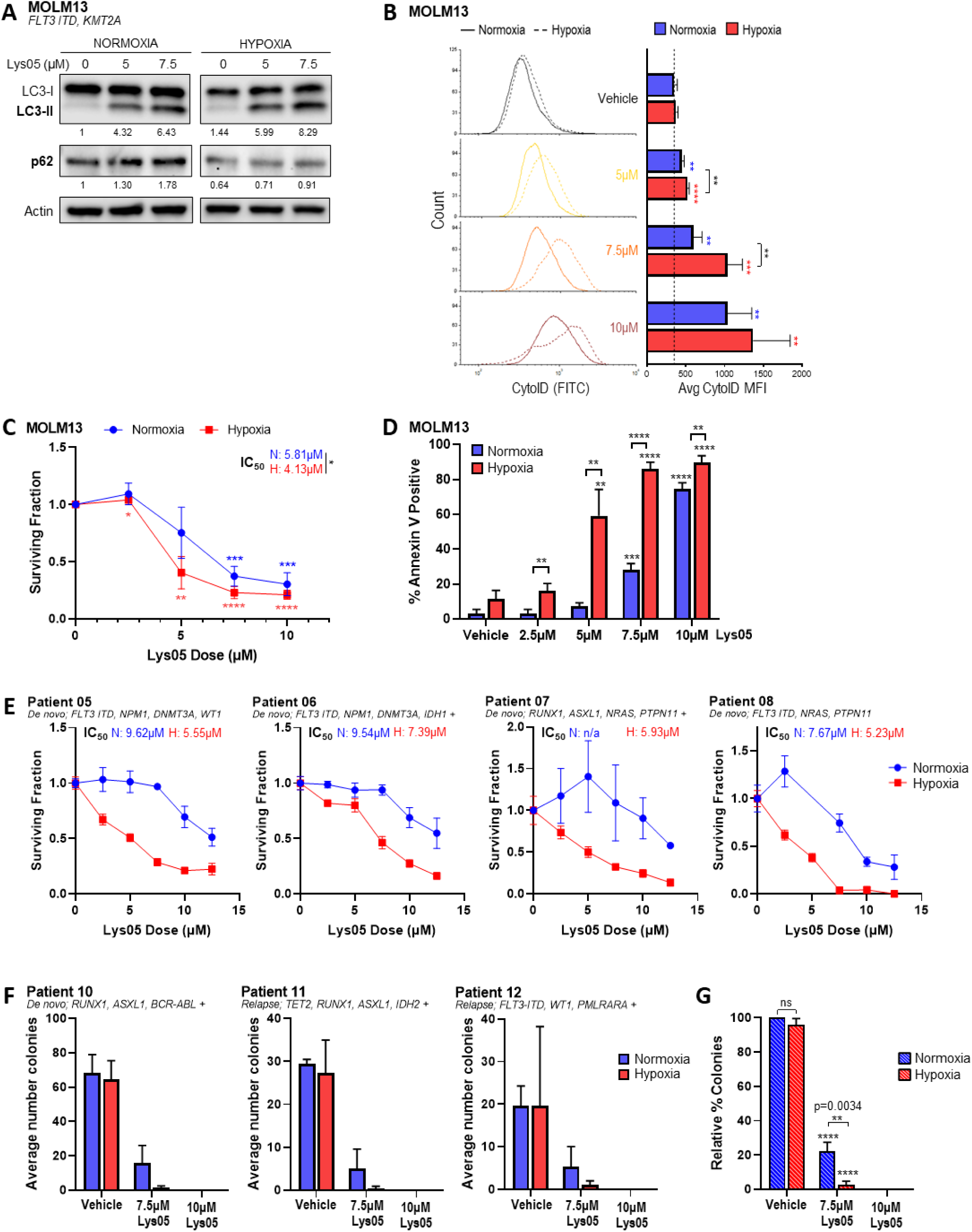
Lys05 exerts enhanced anti-leukemic effects under hypoxia. (A-D) Human AML cell line MOLM13 was cultured under normoxic (21% O_2_) or hypoxic (1% O_2_) conditions with indicated Lys05 dose for 48 hours, unless otherwise indicated. 0µM Lys05 = vehicle (PBS). (A) Immunoblot blot analysis of autophagy markers LC3-II and p62 along with actin loading control in MOLM13 cell line following 48-hour treatment. Quantification shown below respective blot – each marker was normalized to actin then normalized to normoxia vehicle. (B) CytoID flow cytometry for autophagy levels following 72-hour treatment. Quantification shown along with representative CytoID histograms. Solid line indicates normoxia, dashed line indicates hypoxia. (C) WST-8 assay for viability. (D) Annexin V-FITC/PI assay for apoptosis. (E) 48-hour WST-8 assay for viability with primary AML patient bone marrow samples. (F-G) Colony formation assays with AML patient bone marrow in the presence of vehicle (PBS) or Lys05 at the indicated concentrations. Colonies were counted on Day 13. (F) Colony counts for each patient sample. (G) Combined analysis – colony counts for the 3 samples in (A) were normalized to normoxia vehicle. Mutational profile of each cell line or patient sample shown in italics. Stars (*) directly above data points/bars indicate significance compared to vehicle. Normoxia data is compared to normoxia vehicle, hypoxia data is compared to hypoxia vehicle, unless otherwise stated as in (A). (A, B) Representative data of 3 independent experiments shown. (C, D) Average of 3 independent experiments shown. (E, F) Data from single experiment shown due to limited availability of primary samples.

Lys05 more effectively inhibited AML cell growth under hypoxia than normoxia in all cell lines tested (**Figure 2C**, **Table 1, Supplemental Figures 2E-F, 3**). This was reflected by a mean IC_50_ of 5.1µM (range 4.1–7.6µM) under hypoxia as compared with a mean IC_50_ of 8.85µM under normoxia (p=0.0032). There was a corresponding increase in apoptosis levels under hypoxia. In MOLM13 cells, Lys05 (5µM) resulted in 8-fold more apoptosis under hypoxia than normoxia (p=0.0043; **Figure 2D**). Similar results were seen in multiple AML cell lines (**Supplemental Figure 2G-H**). Lys05 also more effectively decreased viability of primary AML patient BM samples with diverse cytogenetic and mutational profiles under hypoxia compared to normoxia (**Supplemental Table 1, Figure 2E**). Together these data suggest that Lys05 more effectively inhibits AML growth under hypoxia due to the increased dependence of leukemia cells on autophagy. We investigated the impact of Lys05 on the growth of leukemic stem and progenitor cells under normoxic vs hypoxic conditions via *in vitro* in colony-formation assays performed with primary patient AML cells from patients with *de novo* and relapsed disease. As shown in **Figures 2GF-G**, Lys05 significantly decreased the number of primary AML colonies in a dose-dependent manner with the greatest activity noted at doses >7.5 µM (p=0.0034) and under hypoxic conditions.

### Lys05 exerts *in vivo* activity in human AML xenograft models

We next evaluated the anti-leukemic activity of Lys05 in our systemic MOLM13-BLIV AML xenograft model. Lys05 treatment decreased leukemia disease burden by approximately 40% – 60% compared to vehicle-treated controls (**Figure 3A-B; Supplemental Table 2).** Lys05 also significantly prolonged median overall survival (p=0.0027; **Figure 3C**) and was relatively well tolerated with no significant differences in weight, white blood cell count, hemoglobin levels, or platelet levels between treatment groups over the duration of the experiment (**Figure 3D; Supplemental Table 2**).

**Figure 3.**
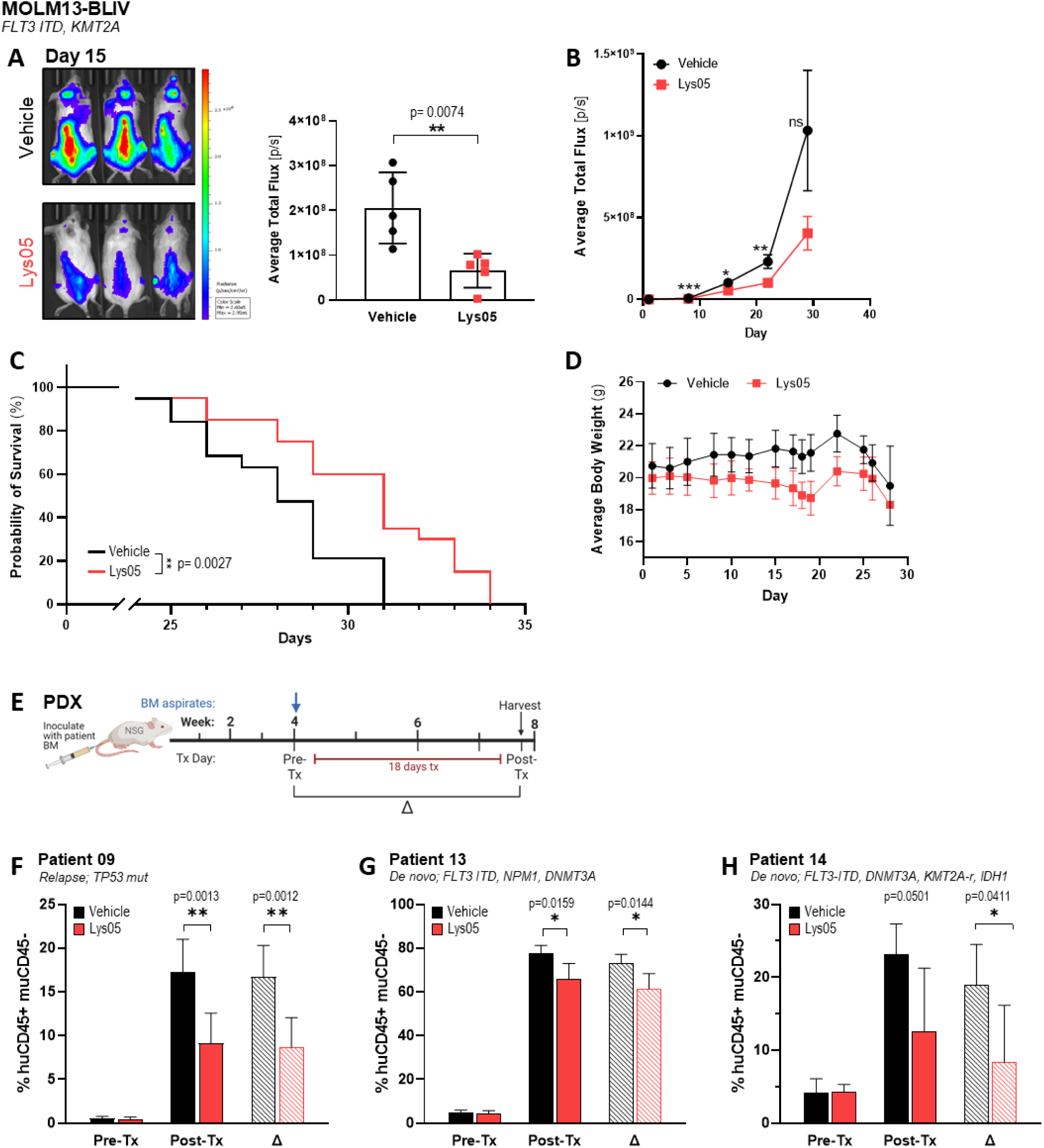
Lys05 exhibits anti-leukemic effects *in vivo*. (A-D) NSG mice were systemically engrafted with luciferase-expressing MOLM13-BLIV human AML cells and treated with vehicle (PBS) or Lys05 i.e., for 18 days. Tumor burden was assessed by BLI. (A) AML tumor burden on day 15 of treatment. Representative BLI images shown (left) with quantification of bioluminescence (right). (B) Leukemia burden over time as measured by BLI. (C) Overall survival. (D) Weights of mice over duration of experiment. (A, D) Individual experiment, n=5. (B, C) Average of 4 independent experiments for a total of n=19. (E-H) NSG mice were inoculated with patient BM cells and engraftment (≥0.1% huCD45+muCD45-in BM) was confirmed by BM aspirates 4 weeks post-inoculation (Pre-Tx). Mice were treated with Lys05 or vehicle (PBS) i.e., for 18 days daily. BM was harvested on day 19 (post-Tx), n=5-8. (E) Experimental design. (F-H) Leukemic burden measured by %huCD45+muCD45- of live cells in BM. Mutational profile of each cell line or patient sample shown in italics. Delta (Δ) = final minus initial (Post-Tx – Pre-Tx); BLI= bioluminescent imaging; BM=bone marrow.

These results were confirmed in three primary PDX AML models. After confirmation of engraftment (≥0.1% huCD45^+^muCD45^-^ in BM) by BM aspirates, mice received treatment with Lys05, or vehicle for 18 days followed by euthanasia and quantification of BM AML disease burden via flow cytometric analysis of huCD45^+^muCD45^--^ cells (**Figure 3E**). In each PDX model, Lys05 significantly reduced leukemia burden (**Figure 3F-H**). In the PDX model established from a patient with *TP53* mutant relapsed AML, Lys05-treated mice had significantly reduced disease burden (9.1% huCD45^+^muCD45^-^ cells) vs. vehicle-treated mice (17.3% huCD45^+^muCD45^-^ cells, p=0.0013; **Figure 3F**). Notably, in this model Lys05 treatment also caused a significant decrease in the CD34+CD38-leukemia stem cell population (p=0.0001; **Supplemental Figure 4**)

### AML cells utilize mitophagy under hypoxic conditions to remove damaged mitochondria

While we have shown that autophagy is critical to acute myeloid leukemia (AML) cell survival within the hypoxic bone marrow microenvironment, it is unknown how autophagy is utilized to provide a survival advantage as autophagy can have many functions within a cell. Based on our previous results, we hypothesized that AML cells rely on mitophagy for survival to remove mitochondria damaged under hypoxia. A larger amount of mitochondrial damage under hypoxia would explain why autophagy inhibitors have enhanced effects under hypoxic conditions. To confirm that mitophagy occurs in AML cell lines, we measured levels of two key mitophagy proteins, PINK1 and BNIP3, under normoxic and hypoxic conditions in the presence of autophagy inhibitors. Since Lys05, CQ, and Bafilomycin (Baf) are late-stage autophagy inhibitors that prevent autophagosome-lysosome fusion, they cause an increase in autophagy (and mitophagy) markers as the autophagosomes and their contents build up due to inhibited degradation. PINK1 initiates the canonical mitophagy pathway and accumulates on mitochondria with depolarized membranes. As shown, Lys05 (and CQ) caused an increase in PINK1 levels under hypoxia alone, while Baf increased PINK1 levels under both normoxic and hypoxic conditions (Figure 4A). Furthermore, Lys05 induced a dose-dependent increase in PINK1 in AML cell lines MOLM13 and HL60, with a further increase in PINK1 under hypoxic conditions (Figure 4B). This is indicative of an accumulation of mitochondria due to lack of clearance by mitophagy and consistent with our previous data demonstrating the distinction between autophagy inhibitors. BNIP3 is involved in the hypoxia-induced mitophagy pathway and is recruited to depolarized mitochondria membranes. Lys05 caused BNIP3 levels to double under hypoxia (Figure 4C). These data indicate that mitophagy is occurring in AML cell lines and is enhanced under hypoxic conditions. It also suggests damage to mitochondria, as indicated by the increased levels of proteins recruited to depolarized mitochondrial membranes.

**Figure 4.**
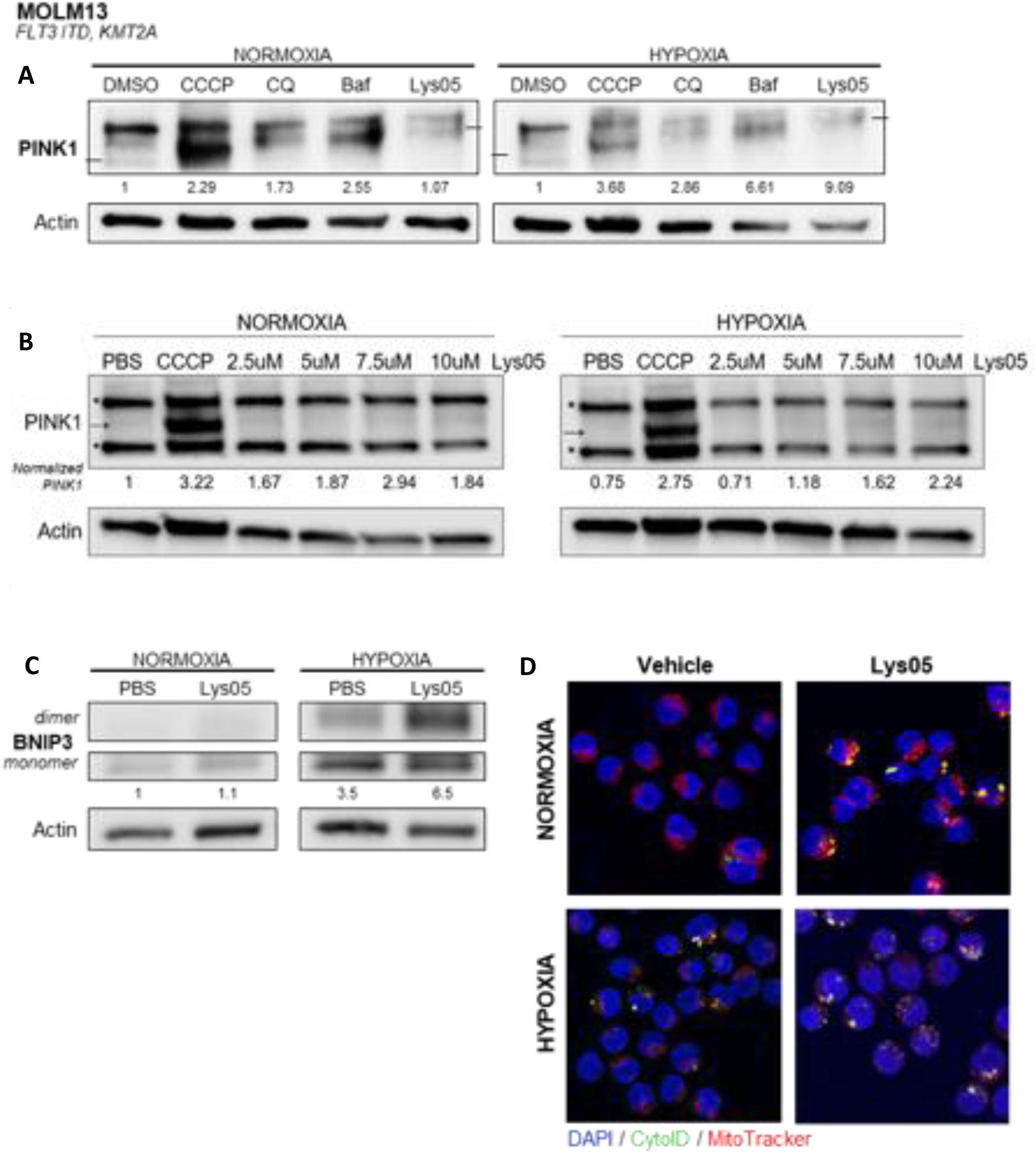
Role of Mitophagy in AML cells under hypoxia. Human AML cell line MOLM13 was cultured under normoxic (21% O_2_) or hypoxic (1% O_2_) conditions with indicated drugs. (A) Immunoblot blot analysis of autophagy marker PINK1 along with actin loading control following 24-hour treatment with vehicle (DMSO), 10µM CCCP, 50μM CQ, 25μM Baf, or 7.5μM Lys05. (B) Immunoblot blot analysis of autophagy marker PINK1 along with actin loading control following 48-hour treatment with vehicle (PBS), 10µM CCCP, or indicated doses of Lys05. Quantification shown below respective blot. Stars indicate non-specific bands. For (A-B), PINK1 was normalized to actin then normalized to normoxic or hypoxic vehicle. CCCP used as a positive control. (C) Immunoblot blot analysis of autophagy marker BNIP3 along with actin loading control following 24-hour treatment with vehicle (PBS) or 5μM Lys05. Total BNIP3 was quantified (dimer + monomer), normalized to actin, then normalized to normoxia vehicle. (D) Fluorescent confocal microscopy images following 24-hour treatment with vehicle (PBS) or 5µM Lys05. Baf= bafilomycin-A1; CQ= chloroquine.

### Mitophagy plays a key role in the anti-leukemic activity of Lys05 under hypoxia

Our observations of the differential effects of late-stage autophagy inhibitors under normoxia and hypoxia along with several recent studies led us to hypothesize that inhibiting autophagy with Lys05 blocks the mitophagy process and prevents the removal of mitochondria, causing an accumulation of damaged mitochondria and ultimately cell death in AML cells. We first compared the anti-leukemic efficacy of late-stage autophagy inhibitors Lys05, CQ, and Baf on human AML cell lines under normoxia and hypoxia. While all three agents inhibit autophagy through deacidification of the lysosome preventing autophagosome-lysosome fusion, their mechanisms of deacidification differ, Lys05 and CQ act through lysosomotropic ion trapping whereas Baf inhibits the vacuole ATPase responsible for pumping hydrogen ions into the lysosome. Furthermore, Baf is known to have additional effects on mitochondria, inducing uncoupling of OXPHOS and depolarization of the mitochondria membrane.^10,11^ Lys05 and CQ demonstrated a dose-dependent increase in apoptosis, with a significant increase in apoptosis under hypoxic conditions, while Baf demonstrated the same level of apoptosis under both conditions (Figure 5A). We hypothesized that these additional mitochondrial effects were responsible for the differences in apoptosis induced. To examine this, we conducted a Seahorse Mito Stress Test following 24-hour treatment with each agent under normoxic or hypoxic conditions. Autophagy inhibition via Lys05 and CQ caused a striking decrease in maximal respiration under hypoxia alone, whereas Baf decreased the maximal respiration rate under both oxygen conditions (Figure 5B). This same difference was also seen in the basal respiration rate, with Lys05 and CQ causing a dose-dependent decrease in basal OCR only under hypoxia whereas Baf caused a stark decrease under both normoxia and hypoxia (Figure 5C).

**Figure 5.**
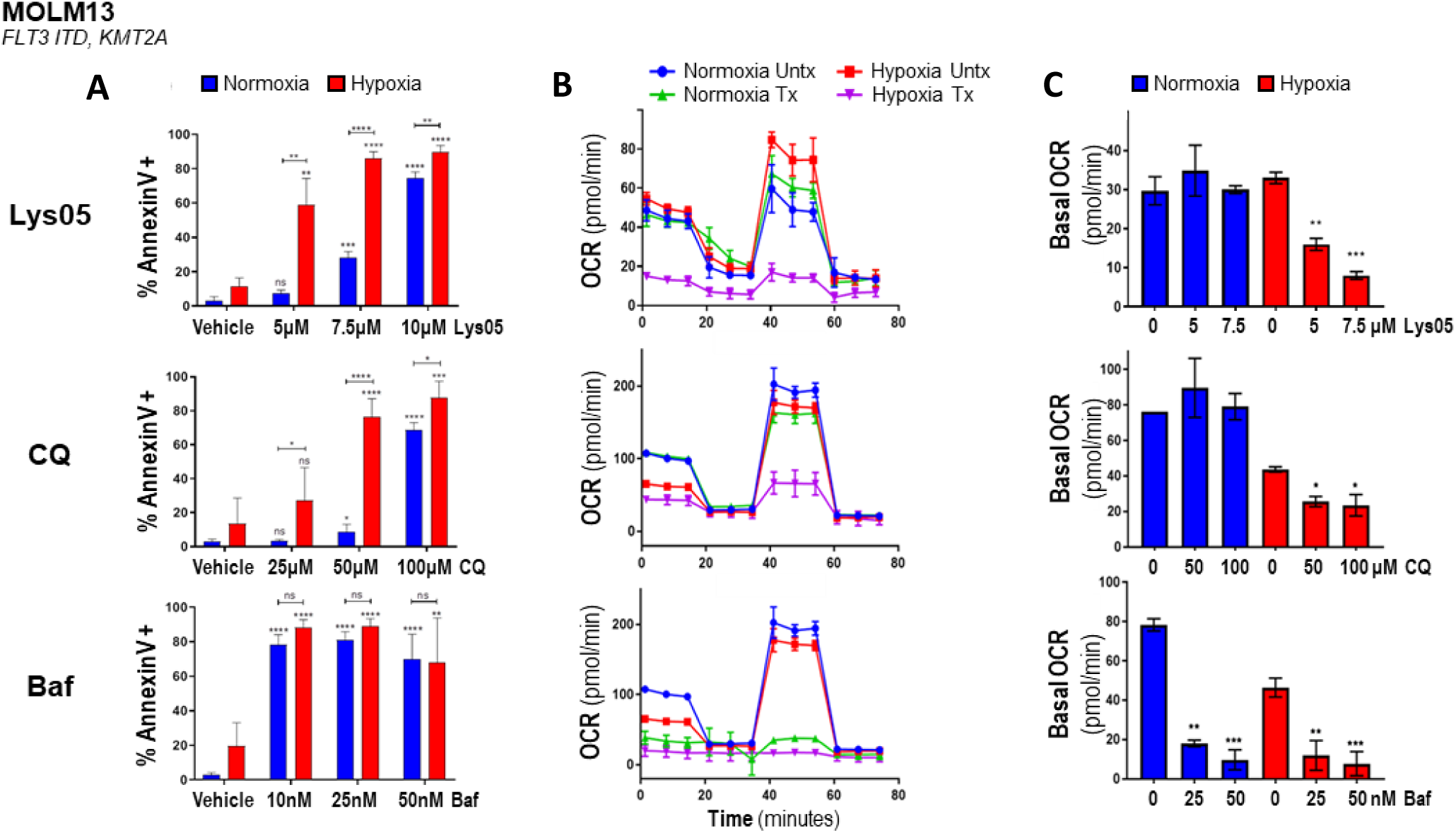
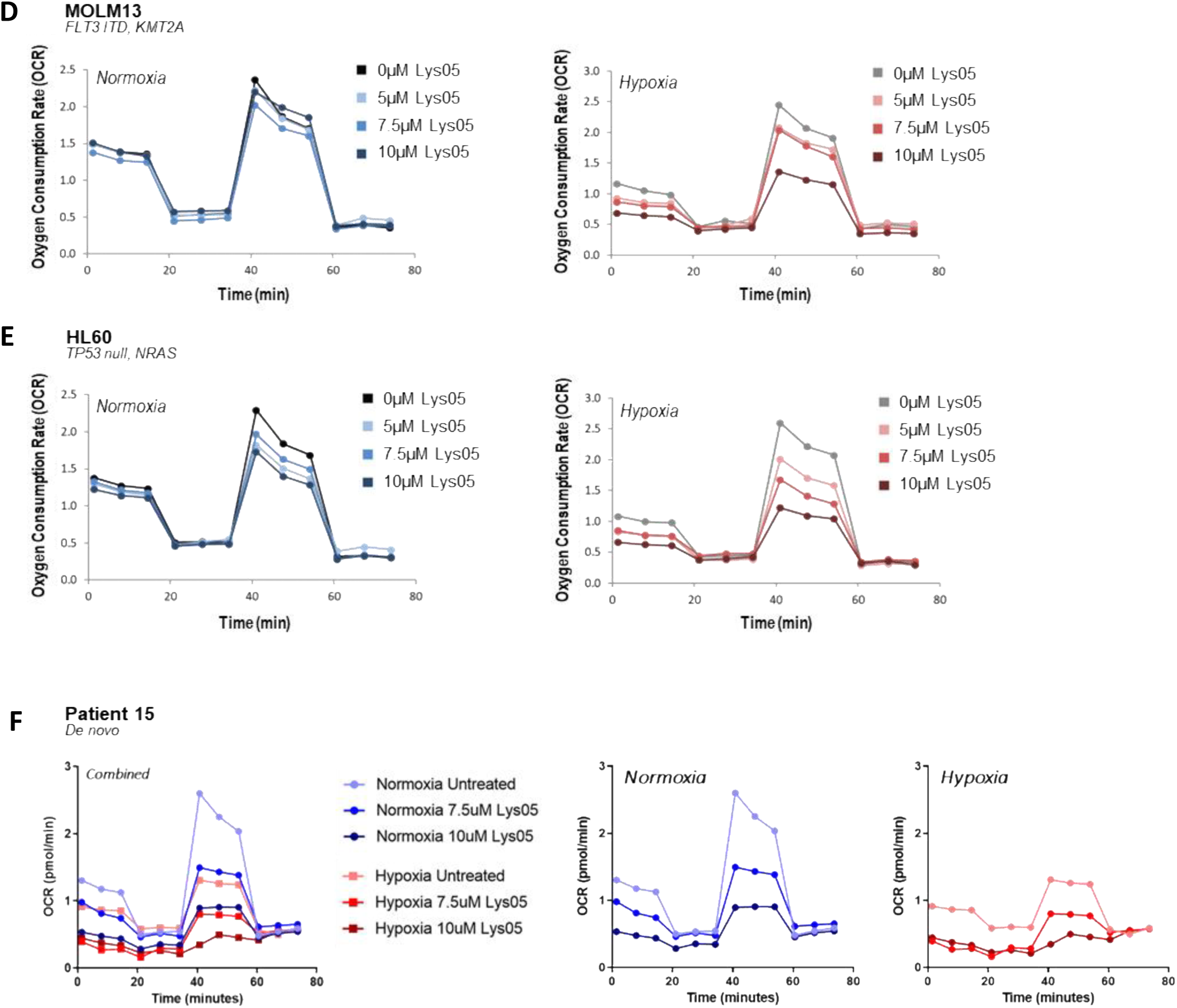
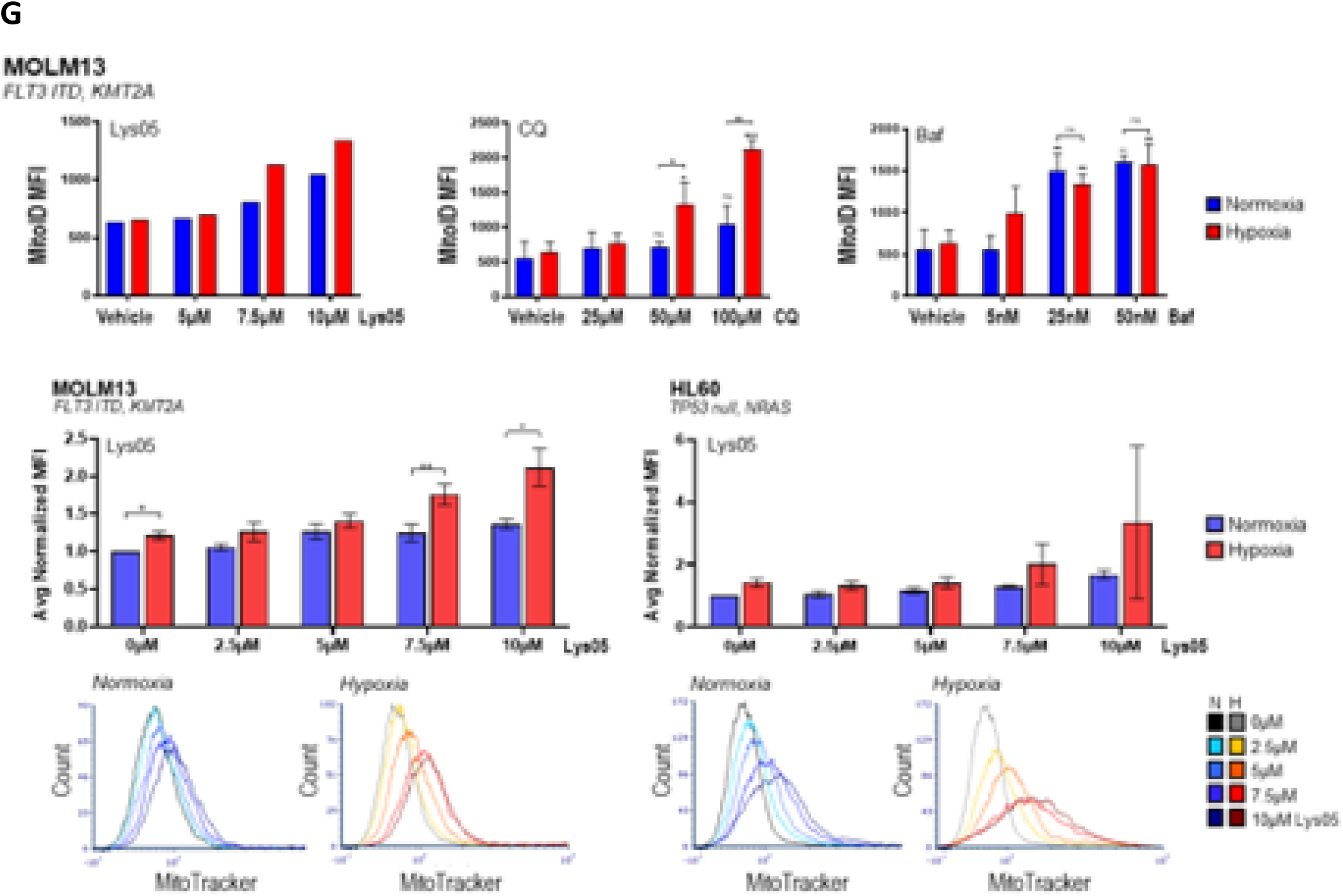
Effects of Lys05 on mitochondrial metabolism and mass under hypoxic conditions. Human AML cell line MOLM13 was treated with Lys05, CQ, or Baf for 24 hours under normoxic (21% O_2_) or hypoxic conditions (1% O_2_) followed by (A) Annexin V/PI flow cytometry for apoptosis, (B, C) Seahorse Mito Stress Test for mitochondrial metabolic rate following treatment with 7.5μM Lys05, 50μM CQ, or 25nM Baf for 48 hours. Stars (*) directly above data points/bars indicate significance compared to vehicle. Normoxia data is compared to normoxia vehicle, hypoxia data is compared to hypoxia vehicle. (D-E) AML cell lines MOLM13 and HL60 were treated with indicated dose of Lys05 for 48 hours followed by Seahorse Mito Stress Test. (F) AML patient bone marrow cells were cultured under normoxic or hypoxic conditions with indicated dose of Lys05 for 24 hours followed by Seahorse Mito Stress Test. (G) Human AML cell lines MOLM13 and HL60 were treated with indicated drug for 48 hours under normoxic (21% O_2_) or hypoxic conditions (1% O_2_) followed by (a) MitoID flow cytometry or (b) MitoTracker flow cytometry for mitochondria mass. (c) Quantification of MitoTracker flow shown (n=3) with representative data below, where darker colors indicate higher doses of Lys05. Baf= bafilomycin-A1; CQ= chloroquine; MFI= mean fluorescence intensity. OCR= oxygen consumption rate; Tx= treated; Untx= untreated.

Furthermore, Lys05 induced a dose-dependent decrease in maximal OCR under normoxic conditions, with a striking further reduction in maximal OCR under hypoxic conditions (Figure 5D). We confirmed these results in an AML patient sample treated for 24 hours with Lys05. As expected, the maximal respiration rate was lower in untreated cells under hypoxia. Treatment with Lys05 resulted in a clear dose-dependent decrease in maximal respiration, with increasing dose of Lys05 further decreasing the respiration rate (Figure 5E-F). Inhibiting autophagy would also block the mitophagic process and prevent the clearance of damaged mitochondria, leading to an accumulation of these organelles and cell death. Indeed, we found that Lys05 and CQ caused a dose-dependent increase in mitochondrial mass under hypoxia alone whereas Baf caused an increase in mass under both oxygen conditions (Figure 5G).^2^

### Lys05 overcomes hypoxia-induced chemoresistance

We further hypothesized that the upregulation of autophagy by leukemia cells in response to treatment accounts for the reduced efficacy of traditional chemotherapy agents in the hypoxic bone marrow, and that this resistance mechanism could be countered with Lys05. We first confirmed that treatment with cytarabine (AraC), venetoclax (Ven), or azacytidine (Aza) resulted in increased autophagy in AML cells as demonstrated by CytoID flow cytometry showing higher levels of autophagic flux under both oxygen conditions (**Figure 6A, Supplemental Figure 5A-B**). We then showed that AML cells are less sensitive (i.e., more resistant) to growth inhibition mediated by each chemotherapeutic agent under hypoxic vs normoxic conditions (**Figure 6B-D, Supplemental Figure 5C-H**). Finally, we determined that Lys05 could overcome this hypoxia-induced chemoresistance by improving the anti-leukemic activity of each agent under hypoxia (**Figure 6B-D, Supplemental Figure 5C-H**).

**Figure 6.**
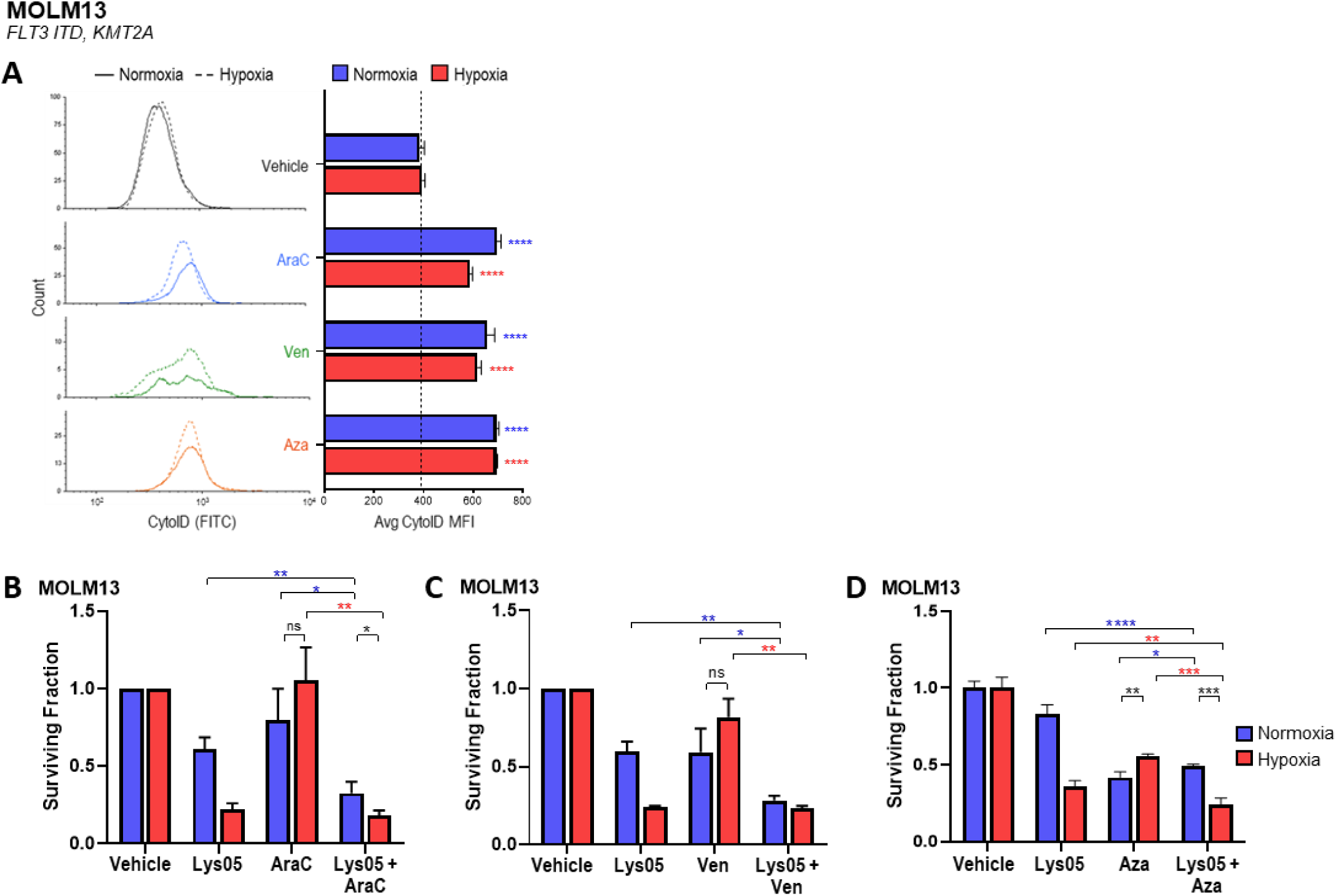
Lys05 overcomes hypoxia-induced chemoresistance *in vitro*. (A) MOLM13 after 72-hour treatment with vehicle (PBS), 0.5µM AraC, 10µM Aza, or 10µM Ven under normoxic (21% O_2_) or hypoxic (1% O_2_) conditions followed by CytoID flow cytometry for autophagy levels. Quantification shown along with representative CytoID histograms. Solid line indicates normoxia, dashed line indicates hypoxia. (B-D) WST-8 assays for viability following treatment of MOLM13 cells under normoxic (21% O_2_) or hypoxic (1% O_2_) conditions. (B) 48-hour treatment with 5µM Lys05, 0.1µM AraC, or Lys05+AraC at the same concentrations. (C) 48-hour treatment with 5µM Lys05, 2.5µM Ven, or Lys05+Ven. (D) 72-hour treatment with 5µM Lys05, 1µM Aza, or Lys05+Aza. All combinations were at the same concentrations as the single agents. Vehicle=PBS. Stars (*) directly above data points/bars indicate significance compared to vehicle. (B, C) Average of 3 independent experiments shown. (A) Representative data of 2 independent experiments shown. (D) Representative data of 3 independent experiments shown. AraC = cytarabine; Aza = azacytidine; MFI= median fluorescence intensity; Ven = venetoclax.

### Lys05 enhances the activity of chemotherapy in AML xenograft models

The addition of Lys05 to AraC was particularly promising, consistently resulting in synergistic anti-leukemic effects over monotherapy across three human AML cell lines under hypoxia (MOLM13 p=0.0022, HL60 p=0.0007, HEL-luc p=0.0028), as demonstrated by combination index values <1 (0.6568 in MOLM13, 0.8191 in HL60, and 0.6916 in HEL-luc; **Supplemental Figure 6A-C**). We therefore tested the combination of Lys05 and AraC *in vivo* in our MOLM13-BLIV xenograft model. The combination significantly decreased leukemic burden over vehicle and both single agent treatments (**Figure 7A-B; Supplemental Table 3**). Combination therapy significantly prolonged survival over AraC (p<0.0001) and vehicle (p<0.0001), with combination-treated mice living 20% longer than AraC-treated mice (**Supplemental Figure 6D**).

**Figure 7.**
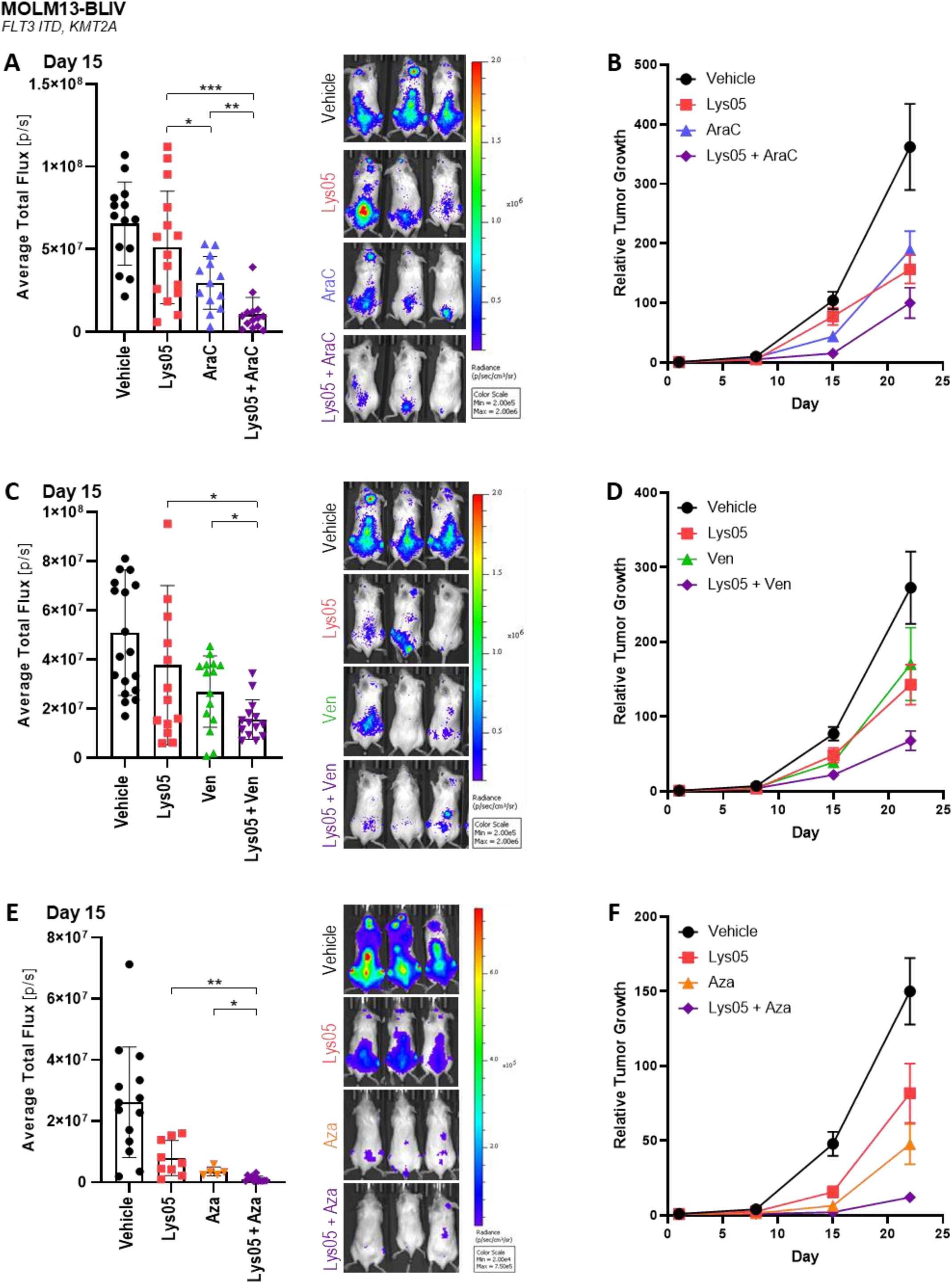
Lys05 overcomes chemoresistance *in vivo*. NSG mice were systemically engrafted with luciferase-expressing MOLM13-BLIV human AML cells. Tumor burden was assessed by BLI. (A, B) Mice were treated with vehicle (PBS), Lys05 (40mg/kg; 18 days), AraC (40mg/kg; 5 days), or combination (same doses). Data shown is an average of 3 independent experiments for a total of n=13-15. (A) AML tumor burden on day 15 of treatment. Quantification of bioluminescence shown (left) with representative BLI images (right). (B) Relative tumor growth over time (each mouse normalized to its day 1 pre-treatment BLI value). (C, D) Mice were treated with vehicle (PBS), Lys05 (40mg/kg; 18 days), Ven (100mg/kg; 5 days on, 2 days off for 3 weeks), or combination (same doses and schedule). Data shown is an average of 3 independent experiments for a total of n=14-19. (C) AML tumor burden on day 15 of treatment. Quantification of bioluminescence shown (left) with representative BLI images (right). (D) Relative tumor growth over time (each mouse normalized to its day 1 pre-treatment BLI value). (E, F) Mice were treated with vehicle (PBS), Lys05 (40mg/kg; 18 days), Aza (2.5mg/kg; 5 days), or combination (same doses and schedule). Data shown is an average of 2 independent experiments for a total of n=5-14. (E) AML tumor burden on day 15 of treatment. Quantification of BLI shown (left) with representative BLI images (right). (F) Relative tumor growth over time (each mouse normalized to its day 1 pre-treatment BLI value). (B, D, F) Error bars represent SEM. AraC = cytarabine; Aza = azacytidine; BLI = bioluminescent imaging; Ven = venetoclax.

To determine whether these effects were limited to cytotoxic chemotherapy alone, we also investigated the ability of Lys05 to improve the *in vivo* efficacy of Ven and Aza in our MOLM13-BLIV model. The combination Lys05+Ven therapy significantly reduced leukemic burden over both single agent Lys05 (p=0.0200) and Ven (p=0.0161) therapy as determined by BLI on treatment day 15 and over time as compared to monotherapy (**Figure 7C-D, Supplemental Table 3**). Similarly, the combination of Lys05 and Aza significantly decreased leukemia burden over single agent Lys05 (p=0.0075) and Aza (p=0.0153) therapy after 15 days of treatment, with this trend continuing over time (**Figure 7E-F, Supplemental Table 3**). There was an overall trend to improved survival in both the Ven and Aza combination therapies (data not shown). These results highlight the therapeutic potential of Lys05 to overcome resistance to frontline chemotherapeutics used against AML.

## DISCUSSION

Growing evidence has highlighted the important role of autophagy in survival, progression, and chemoresistance of acute myeloid leukemia (AML) cells.^16,36^ Upregulation of autophagy genes and signaling pathways has been linked with poor prognosis and refractory disease in AML patients, suggesting that autophagy inhibition could represent a novel therapeutic approach.^37,38^

In AML, the hypoxic marrow microenvironment is believed to contribute to disease relapse by promoting survival of quiescent leukemic progenitor cells within specialized stem cell niches and by rendering AML cells more resistant to cytotoxic agents which are most effective against rapidly dividing cells in an oxygen rich setting. Growth of AML cells under hypoxia also induces environmental stress responses and cellular survival mechanisms, including autophagy.^6,8-11,14^ Leukemic cells residing within a hypoxic marrow microenvironment may be highly dependent on autophagy to survive an inhospitable milieu and therefore could be exquisitely sensitive to potent targeting of autophagy. In our prior work, we showed that AML cells rely on autophagy within the hypoxic bone marrow microenvironment as reflected by the enhanced activity of autophagy inhibitors (chloroquine and bafilomycin) under hypoxic conditions.

Here we provide evidence that AML cells utilize autophagy to clear away mitochondria damaged under hypoxic conditions. Low oxygen conditions have a significant impact on mitochondrial metabolism, leading to enhanced ROS production and mitophagy.^13–15^ Increased mitochondrial damage under hypoxia would necessitate increased mitochondrial clearance via mitophagy. Therefore, inhibiting mitophagy would have a larger impact on hypoxic AML cells, preventing the removal of damaged mitochondria leading to a buildup of these organelles and ultimately cell death.

Currently the only clinically approved autophagy inhibitors are chloroquine (CQ) and its modified form, hydroxychloroquine (HCQ). ^39,40^ CQ is a late-stage autophagy inhibitor that prevents autophagosome-lysosome fusion by accumulating in and deacidifying the lysosome.^34,41^ Prior studies have indicated little to no clinical activity of CQ or HCQ in solid tumor and chronic myeloid leukemia patients treated with CQ alone or in combination with other agents^42,43^ as well as poor *in vivo* pharmacokinetics and lack of marrow penetration.^39,40^ High drug concentrations required to block autophagy in patients are inconsistently achieved *in vivo* and are associated with side effects including arrhythmias, myopathy, nausea and constipation. McAfee and colleagues developed Lys05, a novel modified dimer of CQ that accumulates at higher concentrations within lysosomes, making it a several-fold more potent autophagy inhibitor than CQ and equivalent to genetic knockdown of autophagy in preclinical models.^34,41^ Multiple studies have confirmed that low dose Lys05 exerts improved *in vivo* efficacy over high-dose intermittent dosing of HCQ in solid tumor xenografts.^41,44^ Here we demonstrate that Lys05 exerts several-fold more potent autophagy inhibition than CQ and improved *in vitro* and *in vivo* efficacy over CQ in preclinical human AML models.

Our data show that mitophagy is enhanced in hypoxic AML cells, and that preventing mitochondrial clearance via autophagy inhibition results in increased levels of damaged mitochondria. This indicates that AML cells rely on mitophagy to deal with the increased mitochondrial damage occurring under hypoxic conditions of the bone marrow. We confirm that Lys05 preferentially eradicates human AML cell lines, multiple primary patient samples, and leukemia progenitor cells under chronic hypoxia (1% O_2_). Furthermore, we demonstrate that Lys05 exerts a significant impact on mitochondrial respiration, indicating a link between autophagy and mitochondrial metabolism in AML cells specifically under hypoxic conditions.

Our finding that Lys05 enhances the anti-leukemic activity of multiple AML drugs implicate mitophagy as a key modulator of cell stress and intrinsic adaptive pan-resistance in AML as previously described.^38^ We observed here that the addition of Lys05 consistently enhanced the anti-tumor efficacy of multiple agents (cytarabine, venetoclax, azacytidine) with disparate mechanisms of action in hypoxic AML cells. Moreover, combination Lys05 and cytarabine significantly prolonged survival over cytarabine alone in systemic leukemia xenografts. While not examined here, autophagy upregulation has also been reported to confer resistance of myeloid cells to tyrosine kinase inhibitors, specifically FLT3 inhibitors in AML^45^. ^46^ and JAK inhibitors in myeloproliferative neoplasm.^47^ Inhibition of autophagy improved activity of FLT3 inhibition for FLT3-mutant AML and BCR-ABL inhibition for BCR-ABL positive leukemia in murine models.^23,46,48^ Our data corroborates those of other groups demonstrating targeting mitophagy improves the efficacy of venetoclax in AML cells independent of anti-apoptotic pathways.^49,50^ We show here that Lys05 effectively enhances venetoclax-induced cell death under hypoxia. Other investigators have shown that autophagy inhibition mediated by CQ sensitizes AML cells to MCL1 inhibition in xenograft models, an effect also potentially linked to CQ overcoming *in vivo* hypoxia-induced chemoresistance.^16^

Autophagy inhibition mediated by Lys05 is hypothesized to impact AML survival within a hostile microenvironment and therefore may be effective for treatment of AML subtypes otherwise considered poor prognosis based on underlying disease biology. Here we noted activity of Lys05 across a broad swath of human AML genotypes (including *p53* and *FLT3* mutant disease) as reflected by our panel of multiple human AML cell lines, patient samples, and primary patient derived xenografts. Prior data demonstrating that Lys05 inhibits the growth of *FLT3* mutant (but not wildtype) AML cells was generated in murine leukemia models and may not reflect human AML disease.^23^ Mitophagy is also considered critical for leukemia stem cell biology,^51^ and we show here that Lys05 treatment in vivo reduces both bulk and CD34+CD38-AML cells in a *p53* mutant AML PDX model. Further studies of the efficacy and mechanism of action of Lys05 in genetic knock-down cell lines, additional patient samples and in primary transplantation models are warranted.

While our data supporting the role of mitophagy in hypoxic AML cells is promising, additional experiments to confirm these results would be informative. Examination of mitophagy and mitochondrial damage across different cell lines and patient samples would enhance the rigor of our results. For instance, recent data suggests that SF3B1-mutant AML cells may be particularly susceptible to autophagy inhibition. Genetic manipulation of the mitophagic process via targeted knockouts of mitophagy proteins such as PINK1 to more conclusively show AML cell dependance on mitophagy for survival would be helpful. Moreover, while we have shown that mitophagy occurs in AML cells and that AML cells have damaged mitochondria, further experiments to identify the source of mitochondrial damage and confirm that damaged mitochondria are indeed the cargo of autophagosomes would be informative. Although additional validation of our results using specific pharmacologic inhibitors of mitophagy would be ideal, the existing drugs that interrupt the mitophagic process are non-specific and rely on interfering with mitochondrial fission rather than the mitophagy machinery.

In conclusion, we demonstrate the importance of autophagy (mitophagy) as a critical survival and chemoresistance mechanism of AML cells under hypoxic marrow conditions. Our work showing that AML cells rely on the removal of damaged mitochondria via mitophagy under these conditions identifies a new metabolic vulnerability of AML cells *in vivo*, one that is potentially amenable for future therapeutic exploitation.

## ACKNOWLEDGEMENTS

This research was supported by the Roswell Park Alliance Foundation (Jacquie Hirsch Leukemia Research Fund to ESW) and an NIH National Research Service Award Institutional Training Grant (T32CA085183 to HRSF). This research utilized the Flow and Image Cytometry Shared Resource, Translational Imaging Shared Resource (including the use of the IVIS Spectrum-NIH Shared Instrumentation Grant S10OD16450), Comparative Oncology Shared Resource, and Hematological Procurement Shared Resource at Roswell Park Comprehensive Cancer Center which are supported by National Cancer Institute Cancer Center Support Grant 5P30 CA016056. The authors would like to acknowledge the resource staff for their technical assistance in performing flow cytometric and translational imaging studies, in particular Orla Maguire, Joseph Spernyak, and Steven Turowski. The authors would like to thank Monica Guzman (Weill Cornell Medicine) for provision of cell line MOLM13-BLIV, Michael Becker (University of Rochester) for provision of primary patient sample utilized in our PDX model, along with Pamela Sung and Amanda Przespolewski for their conceptual advice and review of the manuscript.

HRSF is a PhD candidate at Roswell Park Comprehensive Cancer Center. This work is submitted in partial fulfillment of the requirement for the PhD.

## AUTHOR CONTRIBUTIONS

Study conception and experimental design: HRSF, AH, KMD, ESW; Performed experiments: HRSF, MNP, MJ, AH; Analyzed and interpreted data: HRSF, KMD, AH, ESW; Supervised data analysis and interpretation: ESW; Technical support and conceptual advice: MCG; Wrote and edited the paper: HRSF, MCG, ESW.

## SUPPLEMENTAL DATA

### SUPPLEMENTAL METHODS

#### Cell lines and Patient Samples

AML cells were maintained in incubators set with humidity at 37°C, 5% CO_2_, and 21% O_2_ for normoxic conditions or 1% O_2_ for hypoxic conditions using a ProOx C21 Hypoxia Subchamber (BioSpherix, Parish, NY). Cells were cultured in Roswell Park Memorial Institute (RPMI)-1640 Medium or Iscove’s Modified Dulbecco’s Medium (IMDM) medium supplemented with 10% heat-inactivated fetal bovine serum (FBS), 1 U/mL penicillin/ streptomycin, 10mM HEPES, and 1nM sodium pyruvate. All cell lines were monitored for Mycoplasma using the MycoAlert Mycoplasma Detection Kit (Lonza, Basel, Switzerland). Live cells were counted using trypan blue exclusion with a Countess II FL cell counter (Invitrogen, Waltham, MA, USA) or hemocytometer. HEL cells were stably transfected with a pGL4 luciferase reporter vector (Promega, Madison, WI) as described previously^36^ to generate HEL-luc cell line. For experiments, all drugs were added to cells immediately after plating at a concentration of 5×10^5^ cells/mL with continuous exposure, unless otherwise indicated.

#### Human AML cell lines

| Cell Line | Culture Media | Source |
| --- | --- | --- |
| MOLM13 | RPMI-1640 | ATCC |
| MOLM13-BLIV | RPMI-1640 | Monica Guzman, Weill Cornell Medicine (NYC, NY, USA) |
| HL60 | RPMI-1640 | ATCC |
| HEL, HEL-luc | RPMI-1640 | ATCC |
| THP-1 | RPMI-1640 | ATCC |
| U937 | RPMI-1640 | ATCC |
| MV411 | IMDM | ATCC |
| KG-1 | IMDM | ATCC |
| OCI-AML3 | RPMI-1640 | DSMZ |

ATCC (American Tissue Culture Collection), Manassas, VA, USA; DSMZ (Deutsche Sammlung von Mikroorganismen und Zellkulturen), Brunswick, Germany.

### PATIENT SAMPLES

All individuals provided informed consent under Institutional Review Board approved protocols for samples to be stored and used for research purposes. Cryopreserved ficolled mononuclear cells from de-identified patients with AML diagnoses were obtained from institute biorepositories (RPCCC, University of Rochester). Short-term cultures of patient AML cells were performed as previously described.^37^

#### Clinical characteristics of AML patient samples

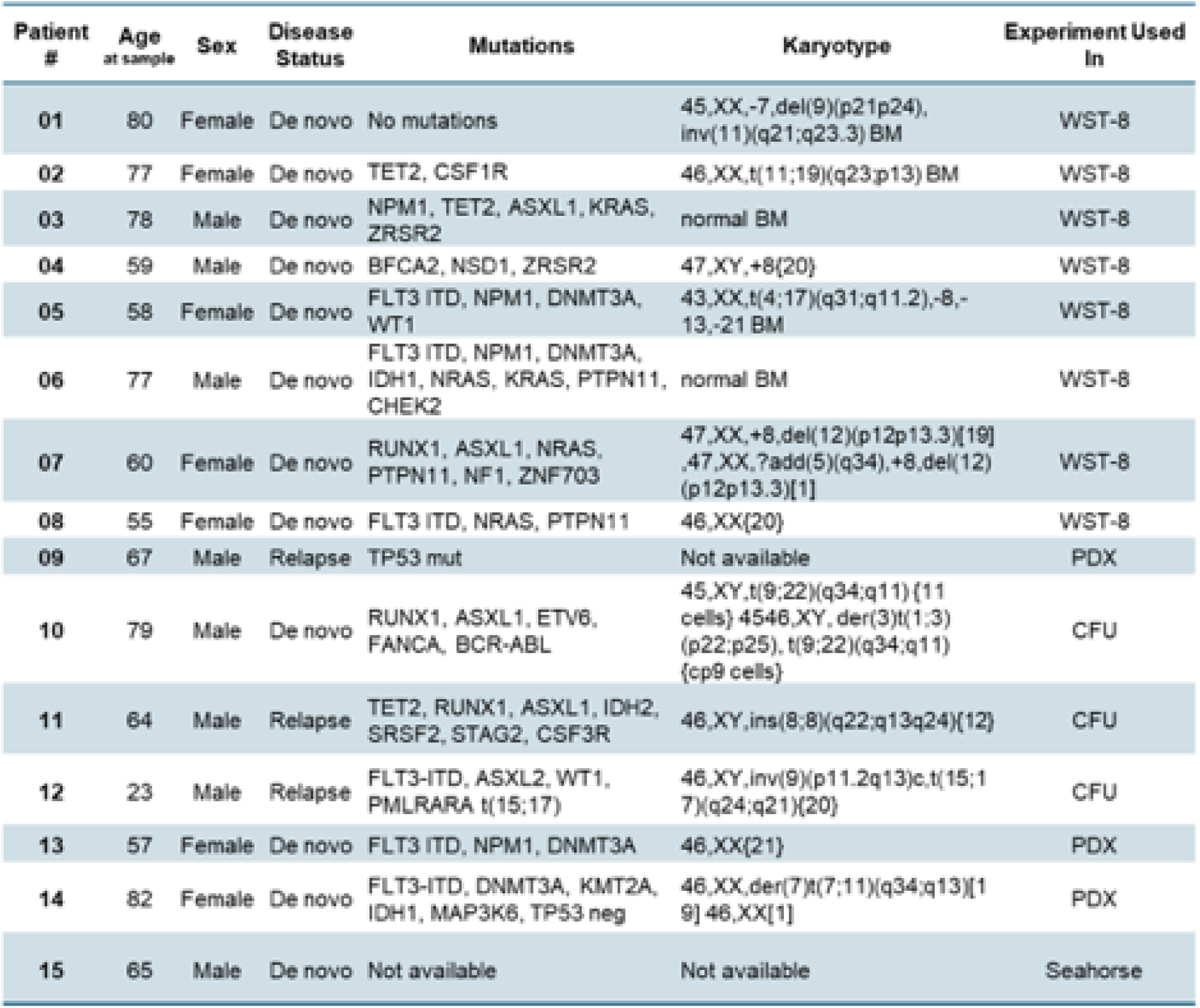

#### Pharmacologic agents

| Drug | Company | Catalog Number |
| --- | --- | --- |
| Lys05 | Selleck Chemicals<br>(Houston, TX, USA) | S8369 |
| Cytarabine (AraC) | Clinical Supply<br>(Roswell Park Comprehensive<br>Cancer Center, Buffalo, NY,<br>USA) | N/A |
| Chloroquine (CQ) | Sigma-Aldrich<br>(St. Louis, MO, USA) | C6628 |
| Bafilomycin (Baf) | Cayman Chemical (Ann Arbor,<br>MI, USA) | 0219391910 |
| Azacitidine (Aza) | Selleck Chemicals | S1782 |
| Venetoclax (Ven) | MedChemExpress (Monmouth<br>Junction, NJ, USA) | HY-15531 |

#### *In Vitro* Assays

For WST-8 (water-soluble tetrazolium salt) assays, absorbance (450nm) was read on an Epoch 2 Absorbance Microplate Reader (BioTek, Winooski, VT).

Flow cytometric analysis was conducted using FCS Express (De Novo Software).

For colony formation assays, drug was dosed to the final volume of MethoCult.

#### Immunoblotting

Cells were lysed in TritonX-100 buffer followed by protein quantification and separation by SDS-PAGE. Triton X-100 buffer: 50mM Tris-HCl, pH 7.4, 150 mM NaCl, 1% Triton X-100, 1% (w/v) supplemented with protease and phosphatase inhibitor cocktail (Sigma-Aldrich, St. Louis, MO, USA). Protein quantification was conducted with a Pierce BCA Protein Assay Kit (Thermo Fisher Scientific, Waltham, MA) and results read with a BioTek Epoch2 Microplate Spectrophotometer (BioTek, Winooski, VT, USA). Equivalent amounts of cell lysate were separated by SDS-PAGE and transferred to PVDF using the Mini-PROTEAN Tetra electrophoresis system (Bio-Rad, Hercules, CA, USA). SDS-PAGE 10-/15-well, 4-20% gradient Bio-Rad Mini-PROTEAN TGX Gels (4561094/4561096). Bio-Rad Precision Plus Protein Standards Kaleidoscope ladder (1610375). Wet transfer was conducted. Detection using HRP-linked anti-mouse/anti-rabbit secondary antibodies (Bio-Rad) and chemiluminescence (ECL plus, GE Healthcare, Pittsburgh, PA). Results were acquired using a LI-COR Imager with Image Studio software (LI-COR) software or Bio-Rad Imager with Image Lab software (Bio-Rad) and quantified relative to actin levels using Image Studio Lite (LI-COR).

#### Immunoblot antibodies

| Antibody | Company | Catalog Number |
| --- | --- | --- |
| Anti-LC3B | Cell Signaling Technology<br>(Danvers, MA, USA) | 2775S |
| Anti-SQSTM1/p62 | Cell Signaling Technology<br>(Danvers, MA, USA) | 8025S |
| Anti-beta Actin (C4) | Santa Cruz Biotechnology<br>(Dallas, TX, USA) | Sc47778 |
| HRP-linked anti-mouse<br>secondary | Bio-Rad<br>(Hercules, CA, USA) | 1706516 |
| HRP-linked anti-rabbit<br>secondary | Bio-Rad<br>(Hercules, CA, USA) | 1706515 |

#### Cell Line Xenograft mouse models

Animal experiments were performed under an Institutional Animal Care and Use Committee approved protocol. Six-eight-week-old NSG (non-obese diabetic/severe combined immunodeficient mice) and NOD.Cg-*Prkdc^scid^ IL2rg^tm1Wjl^*/SzJ (Jackson Laboratories, Bar Harbor, ME) mice were utilized. Treatment began once engraftment was confirmed by bioluminescent imaging (BLI). Engraftment was confirmed by BLI (approximately 7 days post-inoculation: average 5×10^5^ – 1×10^6^ photons/s total flux/mouse). Animals were allocated into treatment groups with similar average leukemia burden (determined by BLI measurements). Mice were clinically monitored daily to thrice per week for signs of morbidity defined as weight loss >20% or hind-limb paralysis, prompting euthanasia. Luciferin (75mg/kg) was injected i.e., two minutes prior to BLI followed by bioluminescent imaging. Vehicle for Lys05, AraC, and CQ was PBS. Vehicle for venetoclax consisted of 10% EtOH, 60% PG50, and 30% PEG400. Vehicle for azacitidine consisted of 5% DMSO, 30% PEG300, and 65% H_2_O.

#### Primary Patient-Derived Xenografts

Primary patient AML derived xenografts (PDX) were established in NSG (non-obese diabetic/severe combined immunodeficient, NOD.Cg-*Prkdc^scid^ IL2rg^tm1Wjl^*/SzJ) mice which were obtained from the Comparative Oncology Shared Resource at Roswell Park Comprehensive Cancer Center in Buffalo, NY. The parent colony for this in-house NSG breeding colony was originally from Jackson Laboratories (Bar Harbor, ME), whose technology transfer office provided approval to the Roswell Park core facility to maintain a breeding colony of mice for research use. To prevent mice bred at Roswell Park from becoming a sub-strain carrying new mutations or other genetic modifications not found in the original Jackson mice, the in-house colony is not isolated from the parent colony by more than ten generations, and the colony is refreshed at regular intervals with stock from Jackson Laboratories.

##### Lys05 treatment in human AML PDX experiments (see Figure 3F-H)

6-8-week-old female NSG mice were inoculated (via tail vein) with 1×10^7^ previously engrafted patient bone marrow (BM) cells, n=10. BM aspirates were conducted at week 4 on all mice followed by flow cytometry to check for engraftment (≥0.1% huCD45+muCD45-). Treatment started shortly after, consisting of 18 days of Lys05 or vehicle (PBS) – 5 days daily (on days 1-5), then every other day up to day 18 (on days 7, 9, 11, 13,15, and 17). On day 19 mice were euthanized and the BM from the femurs and tibias were harvested from each mouse then stained and analyzed by flow cytometry.

##### Staining of murine bone marrow

Samples were ACK lysed to remove red blood cells (ACK lysis buffer: 155mM NH_4_Cl, 10mM KHCO_3_, 10mM EDTA pH 8 in ddH_2_O, pH to 7.2-7.4), counted by trypan blue, and 5×10^5^ cells taken for flow cytometry. Cells were resuspended in Brilliant Stain Buffer (566349, BD Biosciences, Franklin Lakes, NJ), Fc blocked (422302, Human TruStain FcX, Biolegend, San Diego, CA, USA) for 5 minutes, then antibodies added. Samples were stained for 30 minutes at 4°C, then washed and resuspended in FACS buffer (PBS + 2% FBS).

#### Flow cytometry antibodies for PDX antibodies

| **Target** | **Fluorophore** | **Clone** | **Catalog Number**<br>BioLegend |
| --- | --- | --- | --- |
| Mouse CD45 | BV605 | 30-F11 | 103140 |
| Human CD45 | PE-Cy5 | HI30 | 304010 |
| Human CD34 | PE | 581 | 343506 |
| Human CD38 | APC | HB-7 | 356606 |

All acquired from BioLegend (San Diego, CA, USA).

## SUPPLEMENTAL FIGURE LEGENDS

**Supplemental Figure 1.**
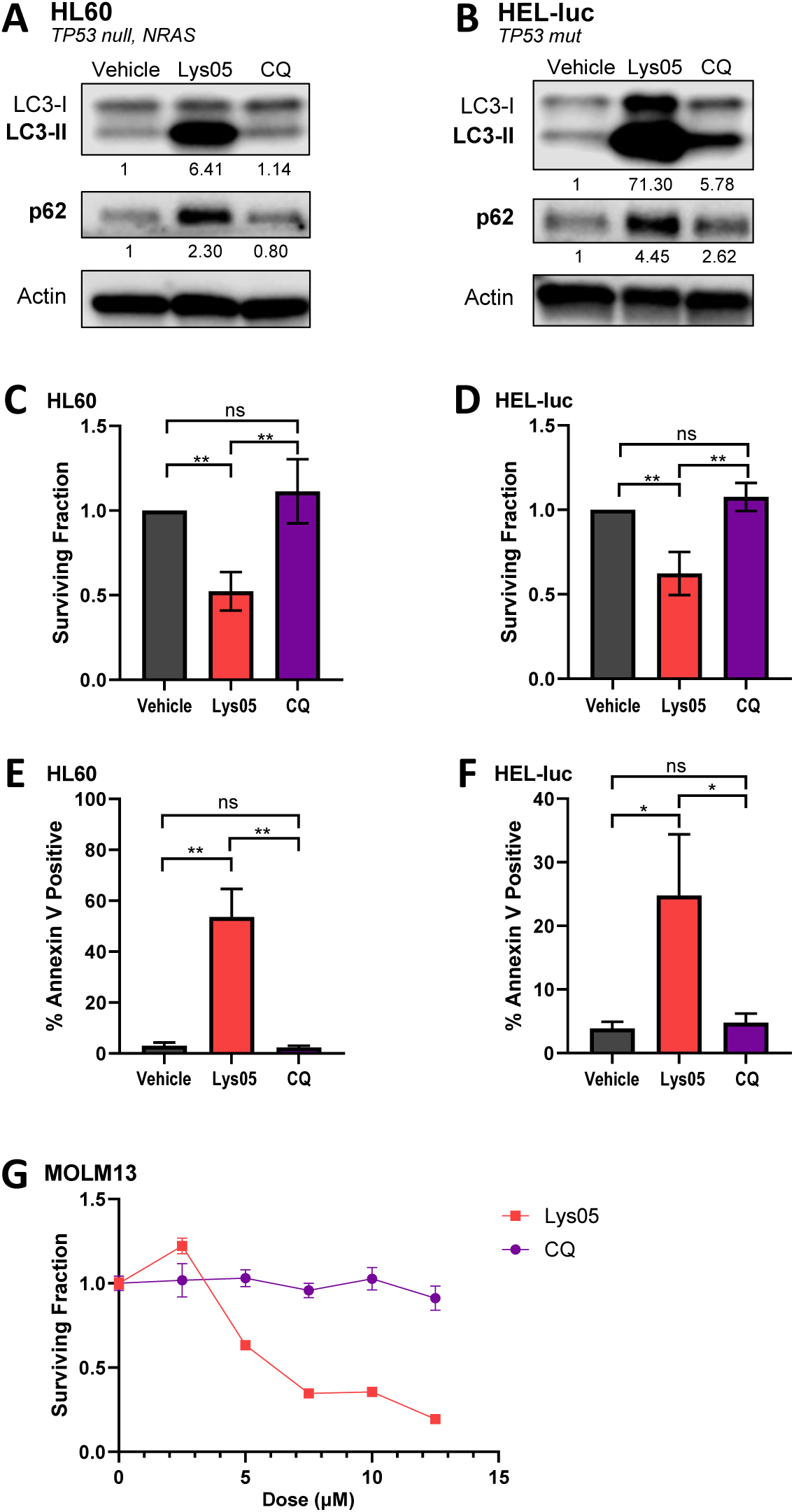
Lys05 exerts greater anti-leukemic activity than CQ in AML cell lines. (A-F) Human AML cell lines HL60 and HEL-luc were treated with vehicle (PBS), 10µM Lys05, or 10µM CQ. (A,B) Immunoblot blot analysis following 48-hour treatment. Autophagy markers LC3-II and p62 shown along with actin loading control. Quantification shown below blot – each marker was normalized to actin then normalized to vehicle. (C,D) WST-8 assay for viability and (E,F) Annexin V-FITC/PI assay for apoptosis following 48-hour treatment. (G) MOLM13 cells were treated with Lys05 or CQ for 48 hours followed by WST-8 assay for viability. Mutational profile of cell line shown in italics. (A,B) Average of 3 independent experiments shown. (C-F) Representative data of 3 independent experiments shown. CQ=chloroquine.

**Supplemental Figure 2.**
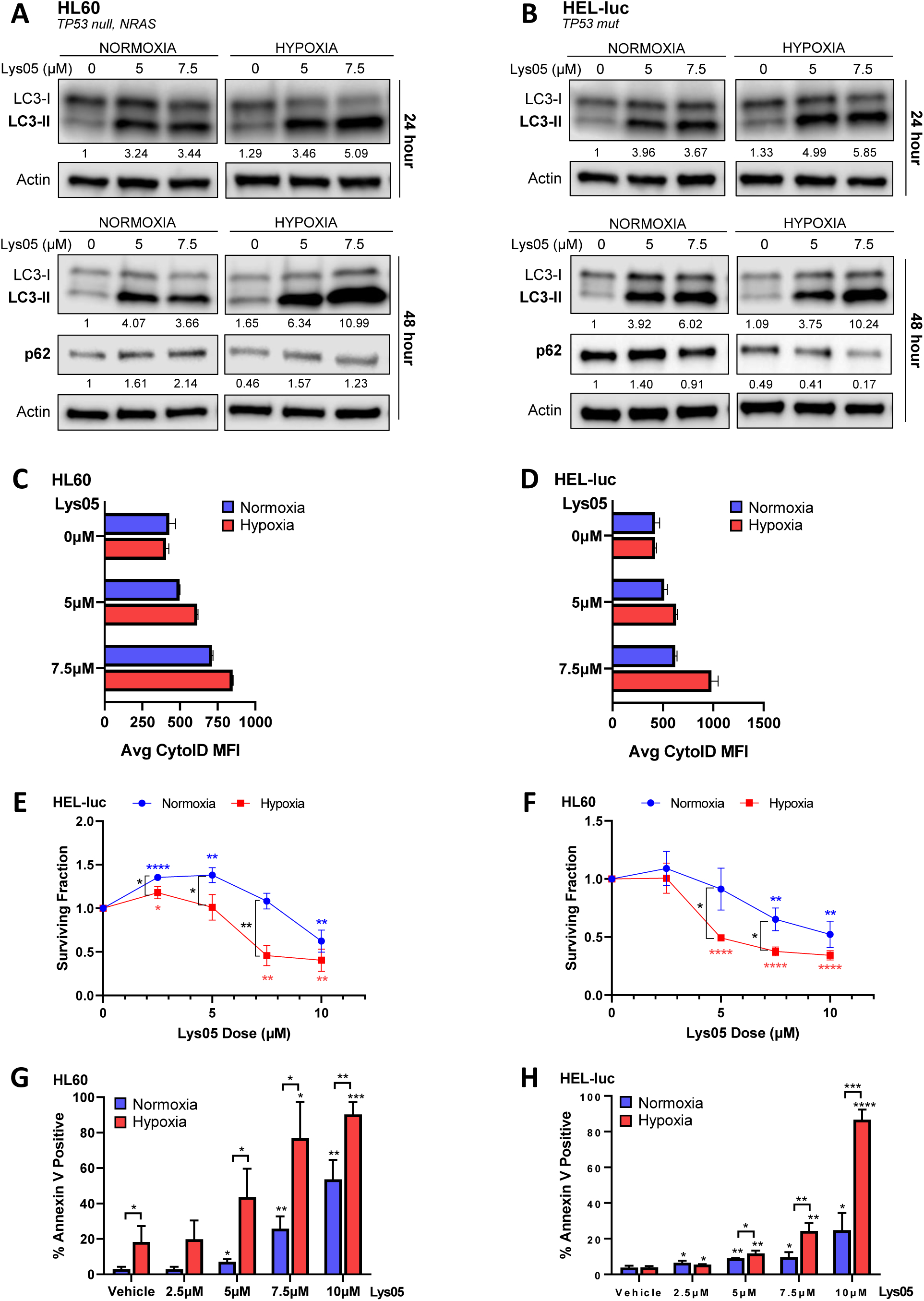
Lys05 exerts enhanced anti-leukemic effects under hypoxia in AML cell lines HL60 and HEL-luc. Human AML cell lines HL60 and HEL-luc were cultured under normoxic (21% O_2_) or hypoxic (1% O_2_) conditions with indicated Lys05 dose for 48 hours, unless otherwise indicated. 0µM Lys05 = vehicle (PBS). (A,B) Immunoblot blot analysis of autophagy markers LC3-II and p62 along with actin loading control in MOLM13 cell line following 24- or 48-hour treatment. Quantification shown below respective blot – each marker was normalized to actin then normalized to normoxia vehicle. (C,D) 72-hour treatment with vehicle (PBS), 5µM, or 7.5µM Lys05 followed by CytoID flow cytometry for autophagy levels. (E,F) WST-8 assay for viability. (G,H) Annexin V-FITC/PI assay for apoptosis. Stars (*) directly above data points/bars indicate significance compared to vehicle. Normoxia data is compared to normoxia vehicle, hypoxia data is compared to hypoxia vehicle, unless otherwise stated as in (A,B). (A-D) Representative data of 3 independent experiments shown (E-H) Average of 3 independent experiments shown.

**Supplemental Figure 3.**
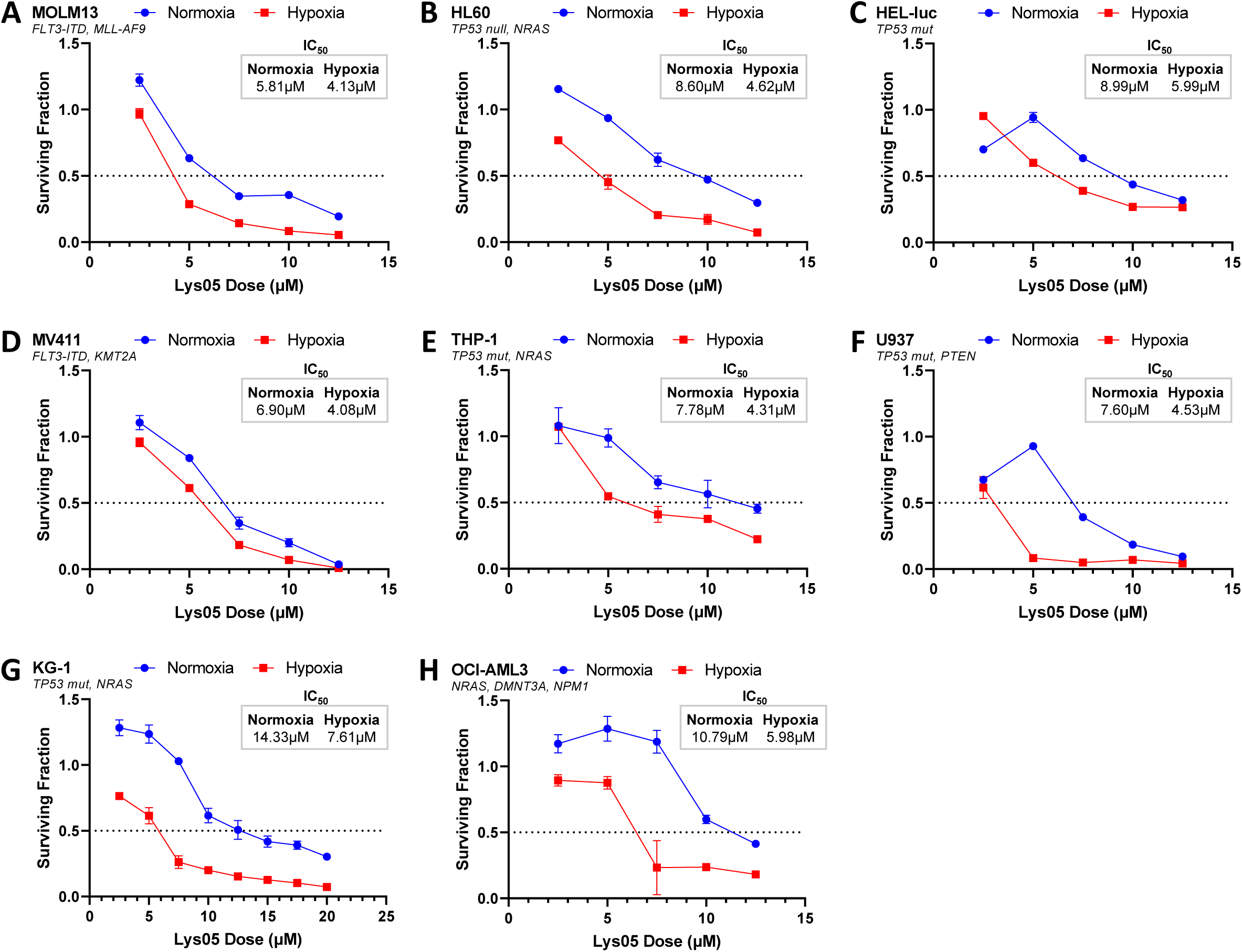
Hypoxia enhances the anti-leukemic effects of Lys05. Human AML cell lines were cultured under normoxic (21% O_2_) or hypoxic (1% O_2_) conditions with indicated Lys05 dose for 48 hours, followed by WST-8 assay for viability. Average normoxic and hypoxic IC_50_ for Lys05 given (n=3-4). Representative data of 3-4 independent experiments shown. Mutational profile of each cell line in italics.

**Supplemental Figure 4.**
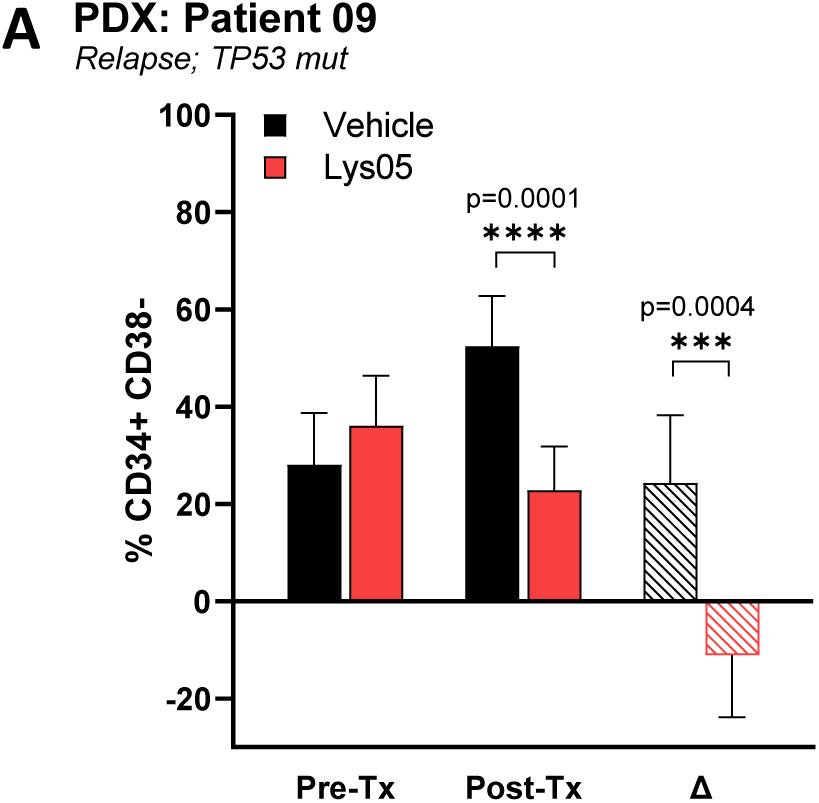
Lys05 reduces the CD34+ CD38-AML cell population in patient-derived xenograft model. NSG mice were inoculated with patient BM cells and engraftment (≥0.1% huCD45+muCD45- in BM) was confirmed by BM aspirates 4 weeks post-inoculation (Pre-Tx). Mice were treated for 18 days with Lys05 or vehicle (PBS) i.p. for 18 days daily. BM was harvested on day 19 (post-Tx). n=5-8. (A) Leukemia stem cell population measured by %CD34+CD38- of live cells in BM. Mutational profile of each cell line or patient sample shown in italics. Delta (Δ) = final minus initial (Post-Tx – Pre-Tx); BM=bone marrow.

**Supplemental Figure 5.**
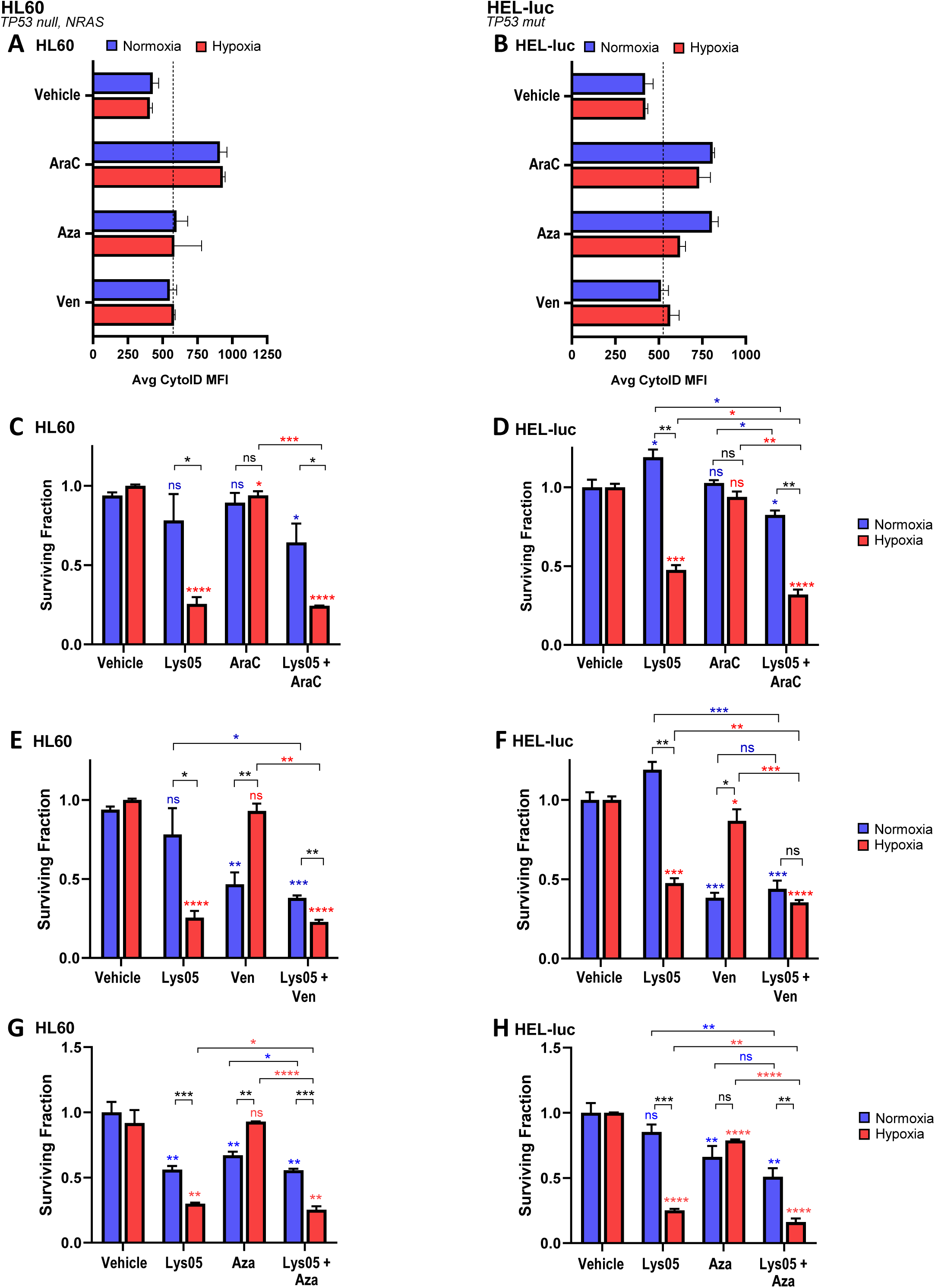
Lys05 overcomes hypoxia-induced chemoresistance in AML cell lines HL60 and HEL-luc. (A,B) Human AML cell lines HL60 and HEL-luc were cultured under normoxic (21% O_2_) or hypoxic (1% O_2_) conditions for 72 hours with vehicle (PBS), 0.5µM AraC (HL60) / 0.25µM AraC (HEL-luc), 10µM Aza, or 10µM Ven followed by CytoID flow cytometry for autophagy levels. Representative data of 2 independent experiments shown. (C-H) WST-8 assays for viability following treatment of HL60 or HEL-luc cell lines under normoxic or hypoxic conditions for 48 hours, unless otherwise specified. Indicated cell line was treated with vehicle (PBS), 5µM Lys05, chemotherapy, or Lys05+chemotherapy at the same concentrations as the single agents. Concentrations for each chemotherapy: (C) 0.1µM AraC (D) 0.01µM AraC (E,F) 2.5µM Ven (G) 1µM Aza (H) 100nM Aza, 72 hours. Stars (*) directly above data points/bars indicate significance compared to vehicle. Normoxia data is compared to normoxia vehicle, hypoxia data is compared to hypoxia vehicle. (C-F) Average of 3 independent experiments shown. (G-H) Representative data of 3 independent experiments shown. AraC = cytarabine; Aza = azacitidine; Ven = venetoclax.

**Supplemental Figure 6.**
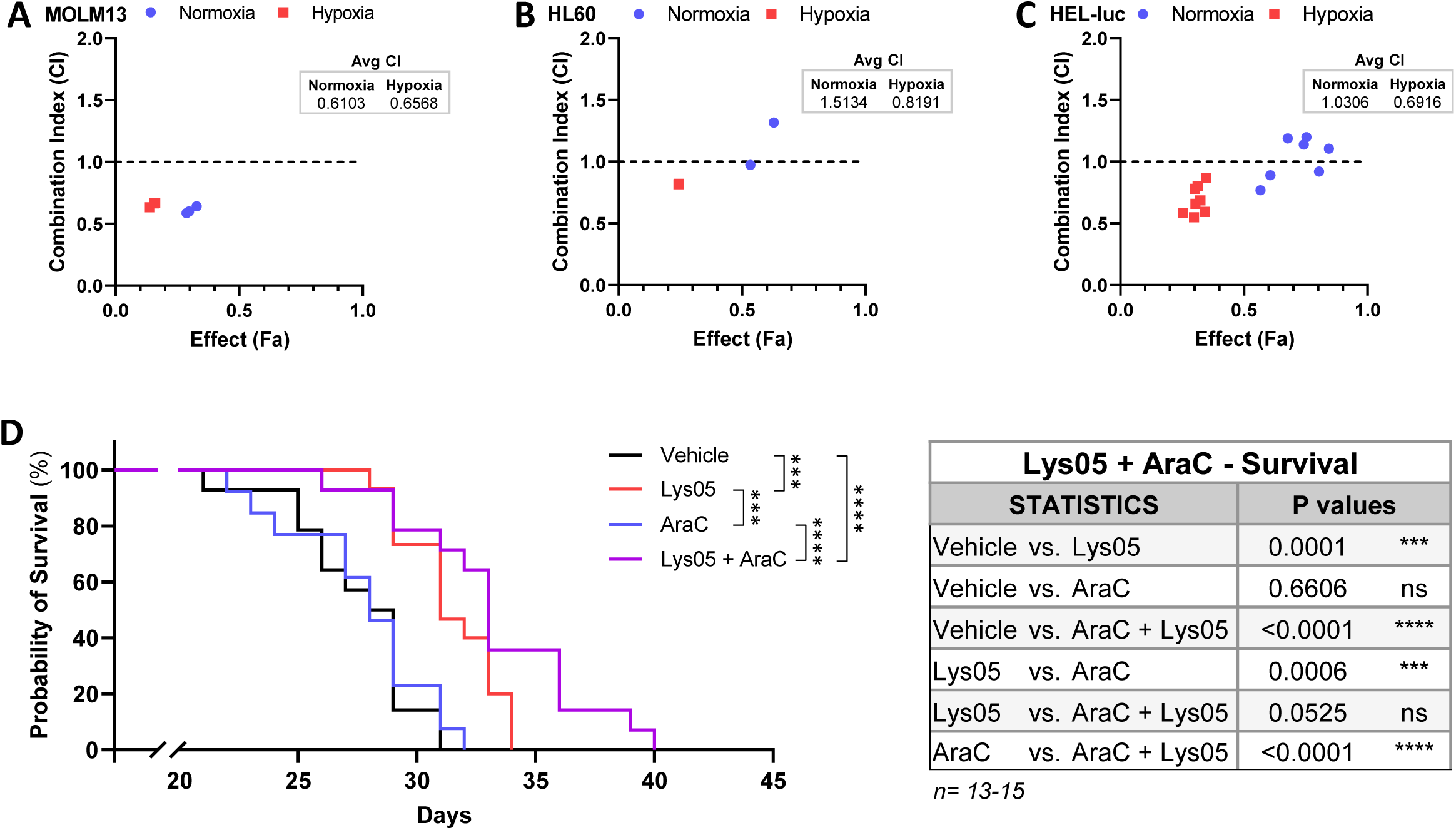
Lys05 enhances the anti-leukemic effects of AraC. AML cell lines were treated with vehicle, 5µM Lys05, 0.1µM AraC (MOLM13/HL60) or 0.01µM AraC (HEL-luc), or Lys05+AraC at the same concentrations as the single agents under normoxic or hypoxic conditions. (A-C) Combination index (CI) plot of Lys05+AraC combination generated using CompuSyn synergy software, CI >1= antagonistic, 1=additive, <1=synergistic. Average CI values for normoxia and hypoxia given. Data from 2-4 independent experiments shown. (D) NSG MOLM13-BLIV xenograft model. Survival curve and statistics following treatment with vehicle, Lys05, AraC, or Lys05+AraC. AraC= cytarabine.

**Supplemental Table 1.** Characteristics of AML patient samples. Age, sex, disease status, mutations present, and karyotype for each patient sample used.

| Patient # | Age<br>at sample | Sex | Disease<br>Status | Mutations | Karyotype |
| --- | --- | --- | --- | --- | --- |
| 01 | 80 | Female | De novo | KMT2A-r | 45,XX,-7,del(9)(p21p24),inv(11)(q21;q23.3) BM |
| 02 | 77 | Female | De novo | KMT2A-r, TET2, CSF1R | 46,XX,t(11;19)(q23;p13) BM |
| 03 | 78 | Male | De novo | NPM1, TET2, ASXL1, KRAS, ZRSR2 | normal BM |
| 04 | 59 | Male | De novo | BFCA2, NSD1, ZRSR2 | 47,XY,+8{20} |
| 05 | 58 | Female | De novo | FLT3 ITD, NPM1, DNMT3A, WT1 | 43,XX,t(4;17)(q31;q11.2),-8,-13,-21 BM |
| 06 | 77 | Male | De novo | FLT3 ITD, NPM1, DNMT3A, IDH1, NRAS, KRAS, PTPN11, CHEK2 | normal BM |
| 07 | 60 | Female | De novo | RUNX1, ASXL1, NRAS, PTPN11, NF1, ZNF703 | 47,XX,+8,del(12)(p12p13.3)[19],<br>47,XX,?add(5)(q34),+8,del(12)(p12p13.3)[1] |
| 08 | 55 | Female | De novo | FLT3 ITD, NRAS, PTPN11 | 46,XX{20} |
| 09 | 67 | Male | Relapse | TP53 mut | Not available |
| 10 | 79 | Male | De novo | RUNX1, ASXL1, ETV6, FANCA, BCR-ABL | 45,XY,t(9;22)(q34;q11) {11 cells}<br>4546,XY,der(3)t(1;3)(p22;p25),t(9;22)(q34;q11) {cp9 cells} |
| 11 | 64 | Male | Relapse | TET2, RUNX1, ASXL1, IDH2, SRSF2, STAG2, CSF3R | 46,XY,ins(8;8)(q22;q13q24){12} |
| 12 | 23 | Male | Relapse | FLT3-ITD, ASXL2, WT1, PMLRARA t(15;17) | 46,XY,inv(9)(p11.2q13)c,t(15;17)(q24;q21){20} |
| 13 | 57 | Female | De novo | FLT3 ITD, NPM1, DNMT3A | 46,XX{21} |
| 14 | 82 | Female | De novo | FLT3-ITD, DNMT3A, KMT2A-r, IDH1, MAP3K6 | 46,XX,der(7)t(7;11)(q34;q13)[19] 46,XX[1] |

**Supplemental Table 2.** Single agent Lys05 in MOLM13-BLIV xenograft model. (Left) Weekly complete blood count data including white blood cells (WBC), hemoglobin (HGB), and platelets (PLT) for vehicle and Lys05-treated mice, along with p-values indicating no significant change in levels with Lys05 treatment, n=5. (Right) Average BLI values (total flux p/s) and statistics for Day 15 of treatment, corresponding to Figure 3A. Weekly quantification of BLI (total flux p/s) and statistics over time, corresponding to Figure 3B. Survival statistics, corresponding to Figure 3C. AraC= cytarabine; Aza= azacitidine; BLI= bioluminescence; Sig= significance; Tx=treatment; Ven= venetoclax.

| Lys05– CBC Data |  |  |  |
| --- | --- | --- | --- |
|  | WBC (10<sup>3</sup>/μl) Avg ± St Dev |  |  |
| Tx Day | Vehicle | Lys05 | P value |
| 1 | 1.77 ± 1.51 | 1.1 ± 0.61 | 0.339 |
| 10 | 2.48 ± 1.56 | 1.63 ± 0.98 | 0.285 |
| 15 | 1.38 ± 0.71 | 0.95 ± 0.69 | 0.310 |
| 22 | 3.88 ± 1.56 | 1.45 ± 0.07 | 0.107 |
|  | HGB (g/dl) Avg ± St Dev |  |  |
| Tx Day | Vehicle | Lys05 | P value |
| 1 | 13.03 ± 0.58 | 13.03 ± 0.26 | 1.000 |
| 10 | 10.85 ± 4.27 | 10.7 ± 4.14 | 0.952 |
| 15 | 9.4 ± 4.97 | 8.73 ± 5.07 | 0.823 |
| 22 | 12.5 ± 0.7 | 7.1 ± 5.94 | 0.109 |
|  | PLT (10<sup>3</sup>/μl) Avg ± St Dev |  |  |
| Tx Day | Vehicle | Lys05 | P value |
| 1 | 435.83 ± 78.07 | 475.5 ± 66.27 | 0.365 |
| 10 | 490.67 ± 240.76 | 595.17 ± 261.26 | 0.488 |
| 15 | 434.33 ± 272.51 | 543.83 ± 322.08 | 0.539 |
| 22 | 473 ± 146.85 | 270.5 ± 275.06 | 0.280 |

| Lys05– Day 15 |  |  |  |
| --- | --- | --- | --- |
| BLI | Average | ± | StDev |
| Vehicle | 2.06E+08 | ± | 7.93E+07 |
| Lys05 | 6.55E+07 | ± | 3.78E+07 |
| STATISTICS |  | P value | Sig |
| Vehicle | vs. Lys05 | 0.0074 | ** |

| Lys05– Timecourse |  |  |  |  |
| --- | --- | --- | --- | --- |
| Tx Day | Avg BLI Vehicle | Avg BLI Lys05 | P value | Sig |
| 1 | 6.80E+05 | 6.76E+05 | 0.9525 | ns |
| 8 | 7.12E+06 | 3.97E+06 | 0.0007 | *** |
| 15 | 1.02E+08 | 5.48E+07 | 0.0159 | * |
| 22 | 2.31E+08 | 1.02E+08 | 0.0086 | ** |
| 29 | 1.03E+09 | 4.05E+08 | 0.0723 | ns |

| Lys05– Survival |  |  |  |
| --- | --- | --- | --- |
| STATISTICS |  | P value | Sig |
| Vehicle | vs. Lys05 | 0.0027 | ** |

Corresponds with Figure 3A
Corresponds with Figure 3B
Corresponds with Figure 3C

**Supplemental Table 3.** Combination of Lys05 and chemotherapy in MOLM13-BLIV xenograft model. Statistics and average BLI values (total flux p/s) for Day 15 of treatment for (Left) Lys05+AraC, corresponding to Figure 5A, (Center) Lys05+Ven, corresponding to Figure 5C, (Right) Lys05+Aza, corresponding to Figure 5E. AraC= cytarabine; Aza= azacitidine; BLI= bioluminescence; StDev= standard deviation; Tx=treatment; Ven= venetoclax.

| Lys05 + AraC – Day 15 |  |  |  | Lys05 + Ven – Day 15 |  |  |  | Lys05 + Aza – Day 15 |  |  |  |
| --- | --- | --- | --- | --- | --- | --- | --- | --- | --- | --- | --- |
| STATISTICS |  | P values |  | STATISTICS |  | P values |  | STATISTICS |  | P values |  |
| Vehicle vs. Lys05 |  | 0.2035 ns |  | Vehicle vs. Lys05 |  | 0.1977 ns |  | Vehicle vs. Lys05 |  | 0.0028 ** |  |
| Vehicle vs. AraC |  | 0.0002 *** |  | Vehicle vs. Ven |  | 0.0027 ** |  | Vehicle vs. Aza |  | 0.0004 *** |  |
| Vehicle vs. AraC + Lys05 |  | <0.0001 **** |  | Vehicle vs. Ven + Lys05 |  | <0.0001 **** |  | Vehicle vs. Aza + Lys05 |  | 0.0002 *** |  |
| Lys05 vs. AraC |  | 0.0413 * |  | Lys05 vs. Ven |  | 0.2526 ns |  | Lys05 vs. Aza |  | 0.0559 ns |  |
| Lys05 vs. AraC + Lys05 |  | 0.0004 *** |  | Lys05 vs. Ven + Lys05 |  | 0.0200 * |  | Lys05 vs. Aza + Lys05 |  | 0.0075 ** |  |
| AraC vs. AraC + Lys05 |  | 0.0015 *** |  | Ven vs. Ven + Lys05 |  | 0.0161 * |  | Aza vs. Aza + Lys05 |  | 0.0153 * |  |
| BLI VALUES | Average | ± | StDev | BLI VALUES | Average | ± | StDev | BLI VALUES | Average | ± | StDev |
| Vehicle | 6.56E+07 | ± | 2.51E+07 | Vehicle | 5.10E+07 | ± | 2.55E+07 | Vehicle | 2.62E+07 | ± | 1.81E+07 |
| Lys05 | 5.12E+07 | ± | 3.41E+07 | Lys05 | 3.78E+07 | ± | 3.24E+07 | Lys05 | 7.93E+06 | ± | 5.76E+06 |
| AraC | 2.97E+07 | ± | 1.60E+07 | Ven | 2.70E+07 | ± | 1.46E+07 | Ven | 3.53E+06 | ± | 1.42E+06 |
| AraC + Lys05 | 1.06E+07 | ± | 1.02E+07 | Ven + Lys05 | 1.57E+07 | ± | 8.04E+06 | Ven + Lys05 | 1.13E+06 | ± | 9.06E+05 |

Corresponds with Figure 5A
Corresponds with Figure 5C
Corresponds with Figure 5E

